# Phylogenomics of *trans*-Andean tetras of the genus *Hyphessobrycon* Durbin 1908 and colonization patterns of Middle America

**DOI:** 10.1101/2022.07.20.500819

**Authors:** Diego J. Elías, Caleb D. McMahan, Fernando Alda, Carlos García-Alzate, Pamela B. Hart, Prosanta Chakrabarty

**Affiliations:** Museum of Natural Science, Department of Biological Sciences, Louisiana State University, Baton Rouge, Louisiana, United States of America; Field Museum of Natural History, Chicago, Illinois, United States of America; Department of Biology, Geology and Environmental Science, University of Tennessee at Chattanooga, Chattanooga, Tennessee, United States of America; SimCenter: Center for Excellence in Applied Computational Science and Engineering, University of Tennessee at Chattanooga, Chattanooga, Tennessee, United States of America; Grupo de Investigación Estudios en Sistemática y Conservación, Universidad del Atlántico-Corporación Universitaria Autónoma del Cauca, Colombia; The University of Oklahoma, Sam Noble Museum of Natural Science, Norman, Oklahoma, United States of America

## Abstract

*Hyphessobrycon* is one of the most species rich and widely distributed genera in the family Characidae, with more than 160 species ranging from Veracruz, Mexico to Argentina. The majority of the diversity of *Hyphessobrycon* shows a *cis*-Andean distribution; only nine species are *trans*-Andean including *H. compressus* (Meek 1908). It is well established that *Hyphessobrycon* is not monophyletic but has been suggested that natural groups can be identified within the larger *Hyphessobrycon* species group. In this study, we test the monophyly of *trans*-Andean species of *Hyphessobrycon* and investigate the placement of *H. compressus*. We have inferred the first phylogenomic hypothesis of *trans*-Andean *Hyphessobrycon* that includes nearly complete taxonomic sampling (eight of nine valid species) using ultraconserved elements (UCEs). We analyzed 75% (1682 UCEs), 90% (1258 UCEs), and 95% (838 UCEs) complete data matrices, and inferred phylogenomic hypotheses under concatenation and coalescent approaches. In all cases, we recovered the monophyly of *trans*-Andean *Hyphessobrycon* inclusive of *H. compressus*, and strong support for three species groups and evidence of cryptic diversity within the widespread *H. compressus* and *H. condotensis*. We used our phylogenomic hypothesis to investigate the biogeographic history of *Hyphessobrycon* in Middle America. Our ancestral range estimation analysis suggests a single event of *cis*- to *trans*-Andean colonization followed by stepwise colonization from the Pacific slope of northwestern South America (Chocó block) to northern Middle America (Maya block). Our work supports the recognition of the *trans*-Andean species as *Hyphessobrycon* sensu stricto and provides a robust evolutionary template to examine morphological characters that will allow us to better understand the diversity of *Hyphessobrycon* in Middle America.

## Introduction

The Neotropics are home to the world’s most diverse assemblage of freshwater fishes, with more than 6200 described species occupying a wide array of aquatic habitats spanning from the Central Mexican Plateau in North America to Tierra del Fuego in South America [1, 2]. The species diversity of this fauna is dominated by ostariophysan lineages: Characiformes (tetras and their relatives), Siluriformes (catfishes), and Gymnotiformes (electric knifefishes), and Acanthomorph lineages: Cichlidae (cichlids) and Cyprinodontiformes (livebearers, killifishes, and their relatives) [2]. Given their high diversity and endemicity, Neotropical freshwater fishes provide exemplar clades to explore the factors that promote the generation of exceptional biodiversity within the region. Such investigations rely on the availability of robust phylogenetic hypotheses for clades of interest, but phylogenetic inference has often been challenged by the rapid diversification experienced by many of these lineages. While efforts to elucidate the evolutionary history of Neotropical freshwater fishes have been vastly benefited from the application of phylogenomic methods [e.g., 3 – 11], there remains considerable uncertainty about phylogenetic relationships and species delimitations, particularly at recent time scales.

Persistent phylogenetic and taxonomic instability is observed within the ostariophysan lineage Characoidei, which includes more than 2150 named species that represent approximately 35% of all Neotropical freshwater fish species [9, 12 – 14]. The characoid genus *Hyphessobrycon* Durbin 1908 (Characidae: Stethaprioninae) is a representative example of this taxonomic uncertainty. This genus is widely distributed from southern Mexico to northern Argentina and includes more than 160 named species [15 – 20] that represent 24% of the species in the hyperdiverse subfamily Stethaprioninae [20]. The majority of *Hyphessobrycon* species (∼95%) are distributed east of the Andes and are hereafter referred to as the *cis*-Andean species. On the other hand, only nine species are distributed west of the Andes (known as the *trans*-Andean species; Fig 1). Four species (*H. ecuadoriensis, H. daguae, H. condotensis*, and *H. columbianus*) are distributed in northwestern South America in the biogeographic Choco, and five species (*H. panamensis, H. savagei, H. bussingi, H. tortuguerae*, and *H. compressus*) are distributed in Central America and southern Mexico (Fig 1).

**Figure 1.**
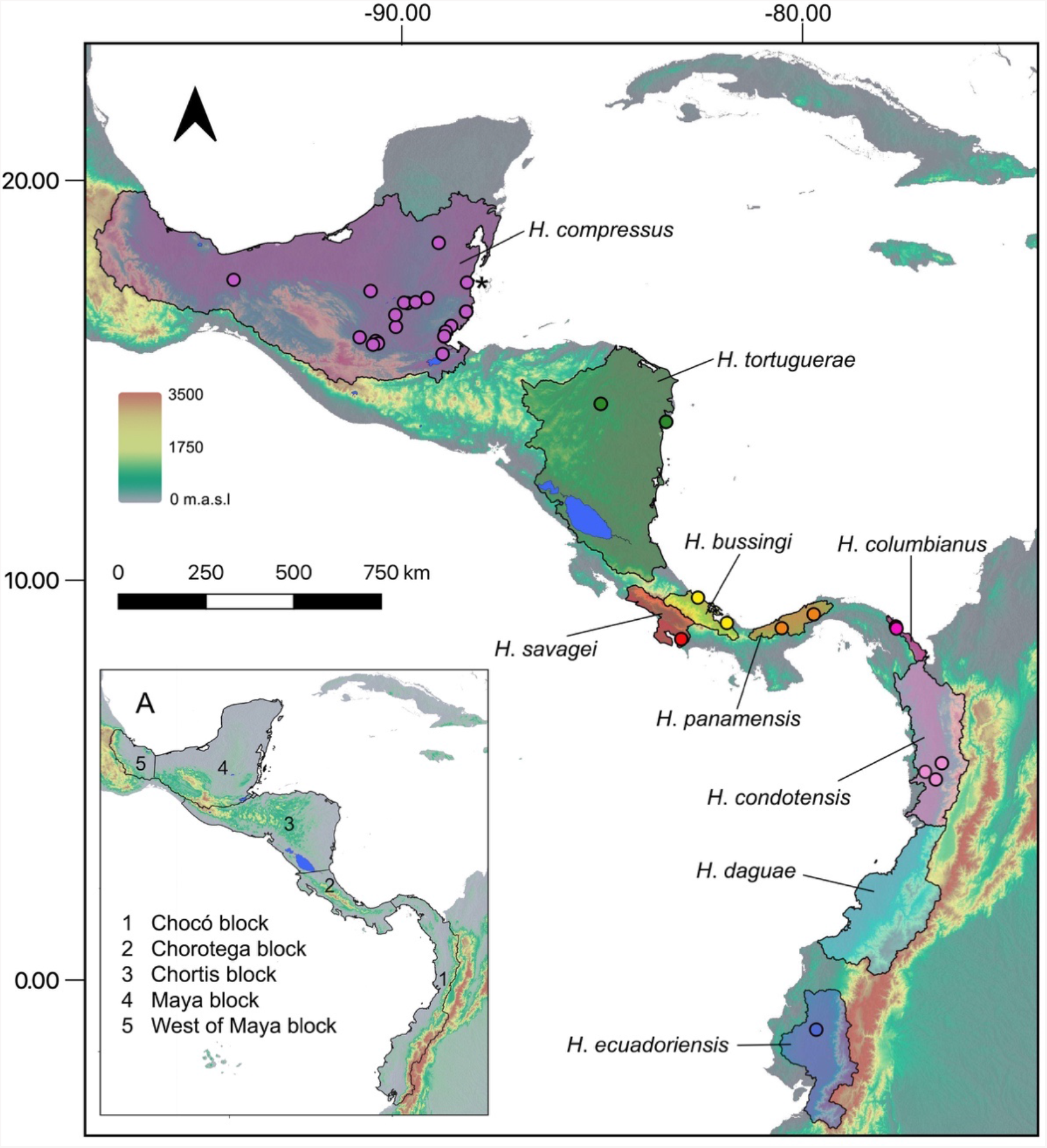
Map of Middle America showing the distribution of the nine valid species of *trans*-Andean *Hyphessobrycon* (shaded regions) based on river basin boundaries (see Materials and methods). Colored circles indicate the geographic location of the samples analyzed in this study. The asterisk indicates the estimated locality of the sample of *H. compressus* from Melo et al. [9] A) Geological blocks in Middle America used as biogeographic units. Km = kilometers, m.a.s.l = meters above sea level.

Previous systematic work based on both morphological and molecular data has shown that *Hyphessobrycon* is not monophyletic [14, 16, 21]. Instead, species currently classified within *Hyphessobrycon* are distributed among multiple different characin lineages, highlighting the need for a phylogenetically-informed taxonomic revision of the genus as currently described [e.g., 12, 14, 22, 23]. Furthermore, the phylogenetic placement of *H. compressus* (Meek, 1904) [24], the type species and most northerly distributed species of *Hyphessobrycon*, is not well understood. Two alternative hypotheses regarding the phylogenetic placement of *H. compressus* have been proposed on the basis of morphological characters and coloration patterns: A) a close relationship to *cis*-Andean species of the ‘rosy tetra’ clade [21, 22] or B) a close relationship with all *trans*-Andean *Hyphessobrycon* [25]. At present, the lack of a robust hypothesis of relationships of *trans*-Andean *Hyphessobrycon* hinders our understanding of the systematics (i.e., which species belong to *Hyphessobrycon sensu stricto*) and the evolutionary history (e.g., biogeographic history) of this group in the northern Neotropics.

Here, we used ultraconserved elements (UCEs) [26], to infer the phylogenomic relationships among *trans*-Andean species of *Hyphessobrycon* under concatenation and coalescent approaches. Specifically, we tested whether *trans*-Andean species of *Hyphessobrycon* are monophyletic, investigated the phylogenetic placement of the type species *H. compressus*, and explored if gene tree heterogeneity present in our dataset provided support for alternative relationships within and among *trans*-Andean species of *Hyphessobrycon*. Furthermore, we evaluated evidence of cryptic diversity within several *trans*-Andean *Hyphessobrycon* species. We then used the resulting phylogeny to infer patterns of colonization of *trans*-Andean *Hyphessobrycon* in the northern Neotropics.

## Materials and Methods

### Taxon Sampling

Our phylogenomic analysis included 95 samples representing eight of the nine valid species of *trans*-Andean *Hyphessobrycon* (Fig 1; S1 Table) and ninePl species of *cis*-Andean *Hyphessobrycon*. Additionally, we included nine characin species as outgroups (S1 Table). Our samples were either collected by the authors or acquired through museum loans, AUM, IMCN, FMNH, LSUMZ, SLU, STRI, UCR, and MZ-UNICACH, and UARC, acronyms follow [27].

### DNA Extraction and Amplification of Mitochondrial Genes

We extracted genomic DNA from a total of 113 samples of characin species using DNeasy spin columns (QIAGEN). We assessed DNA quality and quantity using 1% agarose gels, and a Qubit® 2.0 fluorometer (Invitrogen). To corroborate the taxonomic identity of our samples, we amplified and sequenced the partial Cytochrome Oxidase I (COI) “barcode” gene using the primers and protocol of Ward et al. [28] and compared the sequences against the NCBI database using the nucleotide basic local alignment search tool—blastn—online (https://blast.ncbi.nlm.nih.gov/Blast.cgi) [29].

### UCE Library Preparation

We used ∼300 ng of DNA as starting template and sheared the DNA to a target fragment size of 400 – 600 base pairs (bp) [30] using an EpiSonic Multi-functional bioprocessor (EpicGenTek). We used the Kapa Hyper Prep Kit (Kapa Biosystems) to construct dual index libraries. Libraries were pooled and enriched following protocols in https://www.ultraconserved.org/#protocols with adjustments of Burress et al. [31]. We then used the myBaits UCE Ostariophysan 2.7Kv1 kit (Daicel Arbor Biosciences) to target the capture of 2,708 UCE loci from our samples [32]. Libraries were hybridized and enriched in pools of eight samples. Prior to sequencing, we performed quality control of the final libraries using Bioanalyzer (Agilent). Finally, we combined all libraries in a final pool with a concentration of 10μM, which was then sent for sequencing on one lane of PE150 Illumina HiSeq at Novogene (https://en.novogene.com/). The generated raw reads are archived in the sequence repository archive (SRA) of the National Center for Biotechnology Information (NCBI) (Bioproject PRJNAXXXXX)

We also obtained raw sequences from 87 individuals included in a recently published phylogeny of characoid fishes [9] (SRA BioProject PRJNA563917) that belong to seven sub-families of the family Characidae: Stethaprioninae (63 species), Stevardiinae (six species), Cheirodontinae (three species), Characinae (two species), Exodontinae (two species), Tetragonopterinae (two species), Aphyocharacinae (two species); and Spintherobolinae (four species) (S1 Table). Therefore, our final dataset included a total of 108 species of Characidae, including 20 species of *Hyphessobrycon* representing eight *trans*-Andean species and 12 *cis*-Andean species, including five species assigned to the ‘rosy tetra’ clade sensu Weitzman and Palmer [22].

### Bioinformatics

We used the software PHYLUCE v1.7-1 [33] to perform bioinformatic processing of sequencing reads, identification and extraction of UCE loci from reads, and alignment of UCE loci across samples for downstream phylogenomic inference. Adapter contamination and low-quality reads were removed from our newly generated demultiplexed raw reads using Illumiprocessor (Faircloth 2013), a wrapper for Trimmomatic [34] implemented in the PHYLUCE pipeline using the command *illuminoprocessor*. For the Melo data we used fastp [35] implemented in YAP v0.2.1 (Yet-Another-Pipeline) [36] to perform the cleaning of this dataset. The input file for fastp was generated using the command *new -d* and then we used the command *qc -i* to clean the raw data. Finally, we combined all the clean reads (newly generated + Melo data; FASTA files) into a single folder to continue processing the data.

We assembled the data for each individual sample into contigs using SPAdes [37, 38] within PHYLUCE using the program *phyluce_assembly_assemblo_spades*. To identify and extract UCEs we ran the assembled contigs for all the samples against the ostariophysan probe set (2,708 UCEs) [32] using the program *phyluce_assembly_match_contigs_to_probes*. Then we generated a list of all loci for all samples using the program *phyluce_assembly_get_match_counts* that was used to extract the FASTA data using the program *phyluce*_*get_fastas_from_match_counts*. FASTA files were exploded using the program *phyluce_assembly_explode_get_fastas_file* to obtain summary statistics for each sample assembled. We aligned the UCEs using the MAFFT v.7 aligner [39], first with edge trimming followed by internal trimming using the program *phyluce_seqcap_align*. Subsequently, we trimmed locus alignments using Gblocks [40] using the program *phyluce_align_get_gblocks_trimmed_alignments_from_untrimmed*. Trimmed alignments were cleaned to remove the locus name and just keep the sample name using the program *phyluce_align_remove_locus_name_from_files*. Finally, we generated three data sets that varied in the level of completeness of each alignment (*viz*, 75, 90, and 95%) using *phyluce_align_get_only_loci_with_min_taxa* and calculated summary statistics for each alignment using SEGUL [41]. We used these datasets in downstream phylogenomic analyses. All the data processing steps described above follow standard protocols (https://phyluce.readthedocs.io/) and were implemented on the Louisiana State University High Performance Computing Cluster SuperMIC.

### Phylogenomic Inference

Phylogenomic hypotheses based on the concatenated alignments were inferred for all three datasets (i.e., 75, 90, and 95%) in IQ-TREE 2 [42]. Model selection for each UCE loci was performed using ModelFinder [43] implemented in IQ-TREE 2 followed by tree inference in the same run using the option *-m TEST*. Node support was evaluated using both Ultrafast boostrap (UFBoot2) [43, 44] and the SH-like approximate ratio test (SH-aLRT) [45] with 1000 replicates each. Finally, we estimated gene trees and their support values (UFBoot2; 1000 replicates) for all loci for the three datasets (75, 90, and 95%) in IQ-TREE 2 using the options *-s* and *-bb*.

We inferred a species tree using ASTRAL-III, a two-step method [i.e., inference of gene trees (step 1) and species tree (step 2)] that is consistent with the multispecies coalescent model [46]. ASTRAL-III uses gene trees as input for the inference of the species tree. To minimize the effects of gene tree estimation error, see Simmons and Gatesy [47], prior to inferring the species tree, we collapsed nodes with values <= 50 bootstrap support (i.e., UFBoot2) in all gene trees in Newick utilities [48] using the option *nw_ed b <= 50*. These “collapsed” gene trees were used as input to infer the species tree and to calculate local posterior probabilities (LPP) in ASTRAL-III. In order to have one species or lineage per tip represented in our inferred species tree, we mapped terminal branches of the gene trees representing the same species or lineage into the same taxon (i.e., terminal branch of the species tree) using the option-*a* in Astral-III.

Recently, it has been highlighted that standard measures of support in phylogenomic inference (i.e., bootstrap support and Bayesian posterior probabilities) have some limitations [49, 50]. An expectation is that these measures of support will reach their maximum values even in cases of high gene tree discordance (see Thomson and Brown [50]) possibly leading to high support in topological hypotheses that might not accurately reflect the evolutionary history of the group of interest. We utilized gene concordance factors (gCF) [42, 51] and normalized quartet scores (NQS) [52] as alternative measures of support of relationships in our concatenated and coalescent phylogenomic hypotheses respectively. We used the collapsed gene trees as input to calculate gene concordance factors (gCF), the frequency of the two alternative topologies, (i.e., gene discordance factors; gDF1 and gDF2), and the proportion of gene trees discordant due to polyphyly (gDFP; see Mihn et al.[42]), in our concatenated topology in IQ-TREE 2 using the options --*gcf* and *--df-tree*. We annotated our species tree using the option *-t 16* in ASTRAL-III to calculate the NQS of the main topology (preferred species tree) and its branches and the two alternative topologies. For both of these analyses we focused on the recovered relationships of *trans-*Andean species of *Hyphessobrycon* within Stethaprioninae (monophyly vs non-monophyly). Furthermore, we investigated the recovered relationships between species of *trans-*Andean *Hyphessobrycon*.

### Investigation of Cryptic Diversity

We investigated if there is cryptic diversity within species of *trans-*Andean *Hyphessobrycon* by leveraging *species delimitation* methods that do not rely on prior assignment of samples to species or lineages. Instead of delimiting species our main goal was to use these approaches to explore if there are genetic discontinuities across currently recognized species of *trans*-Andean *Hyphessobrycon*. We generated a reduced UCE dataset (36 samples; S1 Table) with no missing data (i.e., all UCEs are present in all samples) that we analyzed using two different methods: DISSECT/STACEY [53, 54] and SODA [55].

The DISSECT method [53] is implemented in the package “*species tree and classification estimation, Yarely*” (STACEY) [54] in BEAST v.1.8.0 [56, 57] that aims to identify the “minimal number of clusters of individuals” in the dataset [53, 54] as a proxy for the number of species. From our reduced UCE dataset we sub-sampled 100 UCE loci to perform the DISSECT/STACEY analysis (hereafter STACEY). We performed two independent analyses of 500,000,000 generations sampling every 50,000 generations with the following parameters *CollapseHeight*=0.0001, *CollapseWeight*=0.5 with a beta prior (alpha=2, beta=2); *bdcGrowthRate*= lognormal (M=4.6, S=2); *pop-PriorScale* = lognormal (M=-7, S=2), *relativeDeathRate* = Beta (1,1), and *collapseWeight* = Beta (1,1). We evaluated convergence of the analyses using effective sampling size (ESS => 200) for all parameters. We removed 10% of the samples as burn-in and summarized the results of STACEY in the program *SpeciesDelimitationAnalyser* (speciesDA.jar available at http://www.indriid.com/software.html).

We discarded 10% of the sampled trees as burn-in and then we summarized the posterior distribution of sampled trees into a “species or minimal clusters tree” (SMC-Tree) [54] in TreeAnnotator v.1.8.0 [57]. The probabilities of two samples to belong to the same minimal cluster of individuals were visualized in R (version 3.6.1) [58] using the function *plot*.*simmatrix* of the R scrip *plot*.*simmatrix*.*R* (https://github.com/scrameri/smtools/tree/master/SpeciesDelimitation) that generates a probability matrix of assignment of samples and we used the SMC-Tree as a guide to visualize the output of the *plot*.*simmatrix* script.

The ‘*Species bOundary Delimitation using Astral*’ (SODA) [55] is a multi-locus species delimitation method based on quartet frequencies implemented in ASTRAL-III. SODA tests the null hypothesis that a branch has length zero and if the null hypothesis is not rejected, SODA collapses the branch subtending two samples into a “cluster species.” Branches that reject the null hypothesis (i.e., positive branch length) are then used to delimit species [55]. SODA makes two assumptions: 1) there is no gene flow and 2) no population structure; it is worth noting that if these assumptions are violated, SODA can potentially delimit species inaccurately (e.g., over split population-level structure) [55]. We generated gene trees for the reduced dataset in IQTREE-2 and we used the SMC-Tree inferred in BEAST during the STACEY analysis as a guide tree. Both gene trees and the SMC-Tree were used as input for the SODA analysis.

### Biogeographic Analyses

Middle America [59, 60] represents one of the most geologically complex regions on the planet, composed of island-like geological blocks (viz., Maya, Chortis, Chorotega, and Chocó, Fig 1) [60 – 62] that, while presently connected, have undergone multiple episodes of isolation and connectivity at different geological times [63 – 65]. This variation in connectivity among geological blocks likely played a role creating suitable environments at different time periods that allowed the colonization of Middle America from South America by freshwater fishes. Ostariophysan lineages (e.g., tetras and catfishes) [66] were able to colonize Middle America only after the initial closure of the Isthmus of Panama (∼ 20 million years ago) [67 – 69].

We investigated patterns of colonization of *trans*-Andean *Hyphessobrycon* in Middle America [59, 60], using geological blocks (viz., Maya, Chortis, Chorotega, and Chocó, Fig 1) [60 - 62] as biogeographic units. We downloaded occurrence records of *Hyphessobrycon* in Middle America from the global biodiversity information facility (GBIF; www.gbif.org) [70]. We plotted these records after removing those that fell outside the known range of the genus in the region. We generated “distributional” maps for all valid species of *trans*-Andean *Hyphessobrycon* (Fig 1). The species distribution maps were generated by merging major river basins in between the two most distant occurrence records for each species in QGIS 3.4 [71]. River basin boundaries follow those of HydroATLAS [72, 73].

Due to the lack of fossils or geological constraints that will allow us to confidently date our phylogenomic hypothesis, we transformed our maximum likelihood tree (75% complete data matrix) into a chronogram that represents relative divergences time estimates instead of absolute ages. To generate the chronogram, we used the *chronos* function in the R package APE [74, 75]. Then we pruned our chronogram using the function *drop*.*tip* from the package APE [74, 75] to include only one tip per species or lineage of *trans*-Andean *Hyphessobrycon* and the species inferred as their sister South American group in our phylogenomic analyses. The pruned tree was used for downstream biogeographic analyses.

We estimated ancestral ranges in BioGeoBEARS [76] implemented in the software RASP v.4 [77]. We mapped the distribution of each species in each geological block into the pruned tree that we used as input for RASP. *Hyphessobrycon* occurs exclusively in freshwater systems and there is no evidence that it can tolerate salinity. Thus, we assume that dispersal to new areas (biogeographic units) is constrained to existing freshwater connections during geological history. River capture [78] and river anastomosis (e.g., Dias et al. [79]) are the two main processes invoked to explain dispersal (species level) or geo-dispersal (assemblage level) of freshwater taxa across adjacent basins [2]. Therefore, our biogeographic inference was constrained to only allow connections (i.e., dispersal) between adjacent biogeographic units (i.e., geological blocks; Fig 1). Based on the present-day disjunct distributions of all species of *trans*-Andean *Hyphessobrycon* (Fig 1), we constrained our analyses to allow only a maximum of two geological blocks in which one species can be distributed at a time during their evolutionary history. We compared the fit of three different parametric biogeographic models to our data based on the Akaike Information Criterion: Dispersal-Vicariance (DIVA-Like) [76, 80] Dispersal-Extinction-Cladogenesis (DEC-Like) [81] and BAYAREA-Like [82]. All the models were compared with and without the “jump-dispersal” (founder-event speciation) parameter (+J) [76].

## Results

### Alignments summary

#### Phylogenomic inference

We assembled clean reads from 200 samples (see methods) into contigs and identified a total of 2518 UCE loci with a mean length of 443.50 bp (range 37–1926 bp). The 75% complete data matrix contained 1682 UCEs, the 90% complete contained 1258 UCEs, and the 95% complete contained 838 UCEs. After assembly, alignment, and trimming the average alignment length for the 75% complete matrix was 499.11 bp (min = 138, max = 1228 bp) per locus, and the final alignment length was 839,496 bp and contained 337,011 parsimony informative sites (PI) with an average of 200.36 PI per locus. For the 90% complete matrix the average alignment length was 524 bp (min = 145, max = 1118 bp) per locus, and the final alignment length was 659,191 bp and contained 267,932 PI with an average of 212.98 PI per locus. The average alignment length for the 95% complete matrix was 551.39 bp (min = 159, max = 1118 bp) per locus, and the final alignment length was 462,067 bp and contained 187,515 PI with an average of 223.76 PI per locus.

#### Investigation of Cryptic Diversity

For the reduced dataset, we assembled clean reads of 36 samples of *trans*-Andean *Hyphessobrycon* (S1 Table) into contigs and identified a total of 2,235 UCE loci with a mean length of 704.83 bp (range 45 –1691 bp). After assembly, alignment, and trimming the average alignment length for the 100% complete matrix was 794.87 bp (min = 221, max = 1321 bp) per locus, and the final alignment length was 402,998 bp and contained 28,250 parsimony informative sites (PI) with an average of 55.71 PI per locus.

### Phylogenomic Inference

Different levels of missing data did not seem to have an effect on our concatenated phylogenomic inference. Overall, relationships were congruent among the three datasets (i.e., 75%, 90%, and 95% complete matrices; S1 Fig) and the majority of nodes were recovered with high support (UFBoot2 = 100, SH-aLRT = 100). Therefore, we conducted our coalescent, gene concordance factors, normalized quartet scores, and biogeographic analyses based on the gene trees and topology (concatenated and coalescent) inferred using the data matrix with a larger number of loci (75% complete data matrix). We rooted all topologies with the branch subtending to the sub-family Spintherobolinae (Fig 2 and S1 Fig) following previous phylogenomic hypotheses of the family Characidae [9].

**Figure 2.**
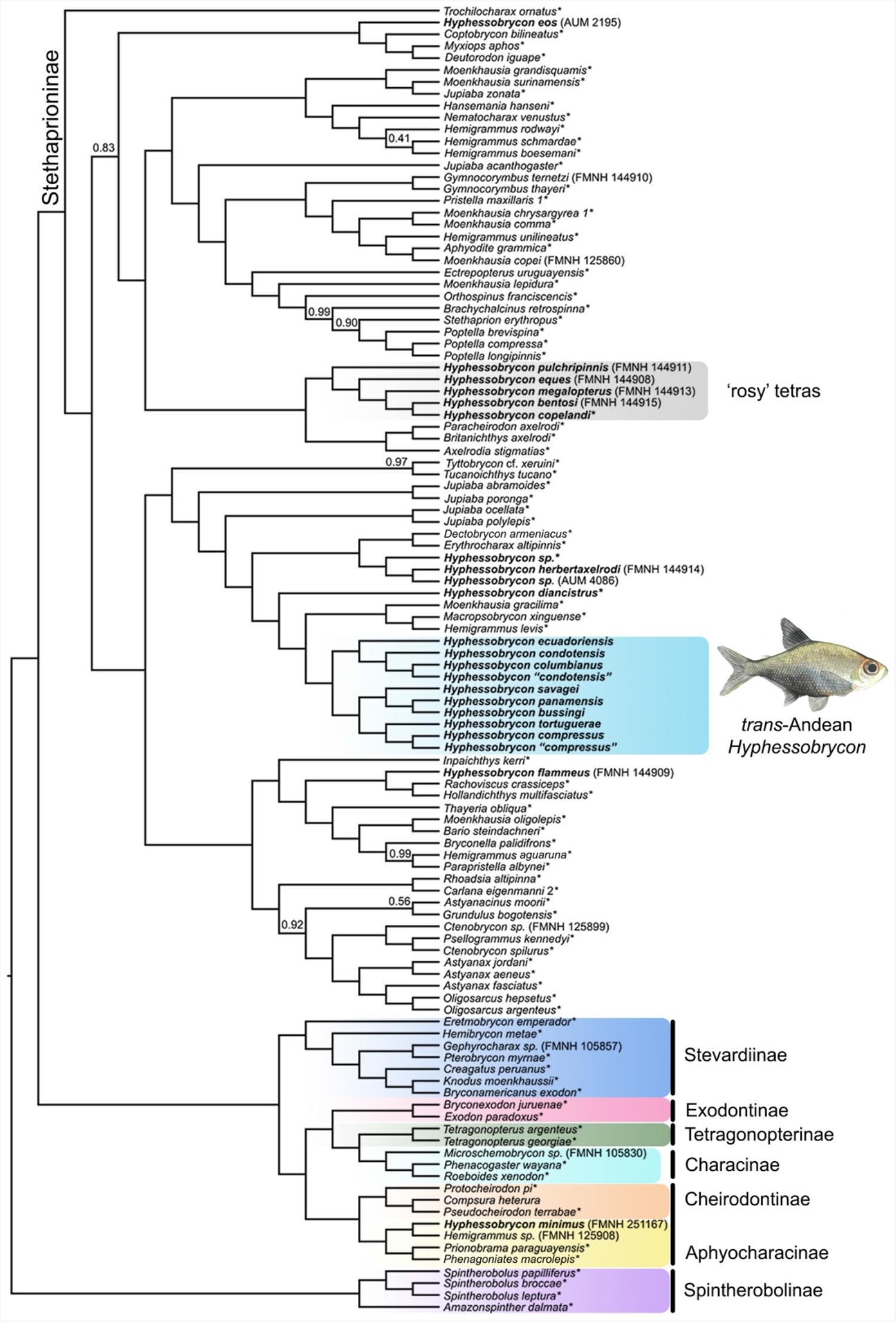
Species tree of the family Characidae inferred with ASTRAL-III based on our 75% complete data matrix (1682 gene trees). All nodes supported by local posterior probabilities (LPP) = 1 unless indicated in the tree. Species of *Hyphessobrycon* are in bold. Asterisks indicate samples from Melo et al. [9]. Illustration of male *Hyphessobrycon compressus* by Duangsamorn Boonwun Boyd.

Overall, the recovered relationships were also congruent between the coalescent and concatenated inferences (Fig 2 and S1 Fig). In all analyses, we recovered the monophyly of the sub-family Stethaprioninae with strong support (LPP = 1, UFBoot2 = 100, SH-aLRT = 100; Fig 2 and S1 Fig), and recovered *Hyphessobrycon* as polyphyletic (n=20; Fig 2 and S1 Fig). The analyzed species of *cis-*Andean *Hyphessobrycon* were scattered among four lineages of Stethaprioninae, some of which were more closely related to *trans-*Andean *Hyphessobrycon* than to other *cis-*Andean taxa. Among the *cis-*Andean *Hyphessobrycon*, we recovered the clade of ‘rosy tetras’ (LPP=1, UFBoot2 = 100, SH-aLRT = 100) but it did not include *H. compressus* (Fig 2 and S1 Fig).

Coalescent and concatenation analyses recovered all the species of *trans*-Andean *Hyphessobrycon* as a monophyletic group, including *H. compressus* (LPP = 1, UFBoot2 = 100, and SH-aLRT = 100; Figs 2, 3 and S1 Fig). The closest relative of the *trans*-Andean *Hyphessobrycon* clade was a clade composed of *Macropsobrycon xinguense, Moenkhausia gracilima*, and *Hemigrammus levis*. Gene concordance factor (gCF) and normalized quartet score (NQS) analyses supported the monophyly of *trans*-Andean *Hyphessobrycon* (gCF= 63.41, NQS = 0.95; Fig 3B) whereas the alternative sets of relationships (i.e., non-monophyly of *trans*-Andean *Hyphessobrycon*) were observed at low frequencies (Fig 3B).

**Figure 3.**
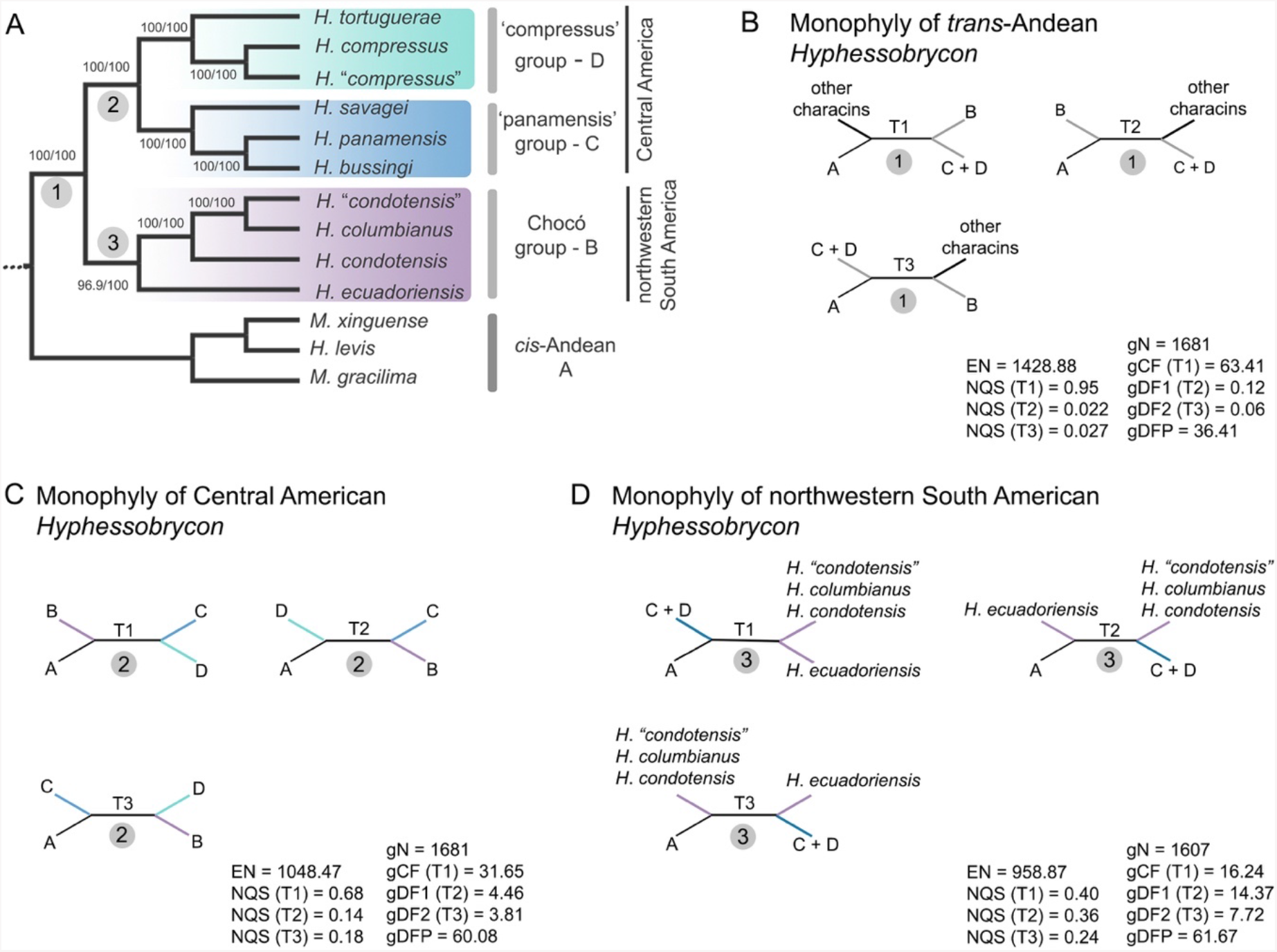
Phylogenomic hypothesis of *trans*-Andean *Hyphessobrycon* and test of alternative sets of relationships in branches of interest. A) Phylogenomic relationships of *trans*-Andean *Hyphessobrycon* inferred using concatenated and coalescent based methods. All nodes supported by local posterior probabilities (LPP) = 1. Support values shown from concatenation analysis, Ultrafast bootstrap (UFBoot2)/ SH-like approximate ratio test (SH-aLRT). Names of species groups and their geographic distribution in Middle America are shown. B-D) Gene concordance factors (gCF) and normalized quartet scores (NQS) for the main topology/quartet (T1) and the two alternatives quartets for a particular branch (T2 & T3): B) monophyly (T1) and non-monophyly (T2 & T3) of *trans*-Andean *Hyphessobrycon*; C) monophyly (T1) and non-monophyly (T2 & T3) of Central American *Hyphessobrycon* (‘panamensis’ and ‘compressus’ species groups); D) monophyly (T1) and non-monophyly (T2 & T3) of northwestern South American *Hyphessobrycon* (Chocó species group). EN = effective number of genes for a branch of interest, NQS = normalized quartet score, gN = number of trees decisive for the branch, gCF = gene concordance factor, gDF1 = gene discordance factor for NNI-1 branch, gDF2 = gene discordance factor for NNI-2 branch, gDFP = gene discordance factor due to polyphyly.

Coalescent and concatenation analyses inferred three well supported groups within the *trans*-Andean *Hyphessobrycon* clade (Fig 3A). The first group included all species distributed in northwestern South America, *H. ecuadorensis, H. condotensis, H. “condotensis*”, and *H. columbianus*, hereafter the Chocó species group (LPP = 1, UFBoot2 = 100, SH-aLRT = 96.9; Fig 3A). Interestingly, samples of *H. condotensis* were not monophyletic (Figs 2 - 3A and S1 Fig), and individuals from the upper Atrato basin (*H. “condotensis”*) were more closely related to *H. columbianus* than to specimens collected from the Baudo and San Juan rivers (*H. condotensis*). The second group was composed of all samples of *H. savagei, H. bussingi*, and *H. panamensis*, hereafter the ‘panamensis’ species group (LPP = 1, UFBoot2 = 100, SH-aLRT = 100; Fig 3A). The third group included two well differentiated lineages of *H. compressus* (LPP = 1, UFBoot2 = 100, SH-aLRT = 100; Figs 2, 3A and S1 Fig): the *H. compressus* lineage from the Usumacinta River in Guatemala and Mexico, the Yucatan Peninsula, and the Coatzacoalcos River and the *H. “compressus”* lineage from aquatic systems in the Izabal lake basin in Guatemala, southern Belize to northern Belize (LPP = 1, UFBoot2 = 100, SH-aLRT = 100; Figs 1 - 3A and S1 Fig), and *H. tortuguerae*, hereafter the ‘compressus’ species group (LPP = 1, UFBoot2 = 100, SH-aLRT = 100; Fig 3A). The Chocó species group was recovered as sister to a clade composed of the ‘panamensis’ and ‘compressus’ species groups: (Chocó, (‘panamensis’, ‘compressus’)). This relationship was congruently recovered by all inference methods (Figs 2 - 3 and S1 Fig) and further supported by gCF = 31.65 and NQS = 0.68 (Fig 3C). The alternative relationships among species groups: (‘compressus’, (‘panamensis’, Chocó)) or (‘panamensis’ (‘compressus’, Chocó)), were observed at relatively low frequencies when compared with the main topology, gCF = 4.46, NQS = 0.14, and gCF = 3.81, NQS = 0.18 respectively (Fig 3C).

Within the Chocó species group, *H. ecuadoriensis* was sister to a clade composed of *H. condotensis, H*. “*condotensis*”, and *H. columbianus* with high support (LPP = 1, UFBoot2 = 100, and SH-aLRT = 96.9), where *H. condotensis* was sister to *H*. “*condotensis*” + *H. columbianus*; ((*H. ecuadoriensis*, (*H. condotensis*, (*H*. “*condotensis*”, *H. columbianus*))) (Fig 3A). Despite the high support observed in both inference methods, the placement of *H. ecuadoriensis* was not unambiguously resolved. The gCF and QNS for the monophyly of the Chocó species group (T1; Figs 3A and 3D) and the alternative topologies (non-monophyly of the Chocó species group; T2 and T3, Fig 3D) were recovered in roughly similar proportions. For the monophyly of the Chocó species group (T1; Fig 3D), gCF=16.24 and NQS=0.40, and for the alternative topologies (i.e., non-monophyly of the Chocó species group) gCF were 14.37 and 7.72, and NQS were 0.36 and 0.24 for T2 and T3, respectively (Fig 3D).

Within the ‘panamensis’ species group, both analyses of UCEs inferred *H. savagei* sister to *H. bussingi* + *H. panamensis*, with high support (LPP = 1, UFBoot2 = 100, SH-aLRT = 100; Fig 3A). In the ‘compressus’ species group, *H. tortuguerae* was sister to *H. compressus* + *H*. “*compressus*”, with strong support (LPP = 1, UFBoot2 = 100, SH-aLRT = 100; Fig 3A).

### Investigation of Cryptic Diversity

The species tree inferred for the reduced dataset of 36 samples and no missing data supported the same topology of *trans*-Andean *Hyphessobrycon* species as the concatenated and coalescent trees of the complete data set (Figs 2 and S1 Fig). Both species delimitation methods uncovered cryptic diversity within *trans*-Andean *Hyphessobrycon* (Fig 4). In the STACEY analysis, we investigated alternative clustering schemes after summarizing 70% of the distribution of post-burnin sampled trees. We identified three clustering schemes: the “optimal” clustering scheme recovered nine lineages within the *trans*-Andean *Hyphessobrycon* and was observed in 48% of the sampled trees, the second clustering scheme recovered 10 lineages and was observed in 15% of the sampled trees, and the third clustering scheme recovered 11 lineages and was observed in 9% of the sampled trees (Fig 4). In contrast, in the SODA analysis, we identified 15 lineages within the samples of *trans*-Andean *Hyphessobrycon* analyzed (Fig 4).

**Figure 4.**
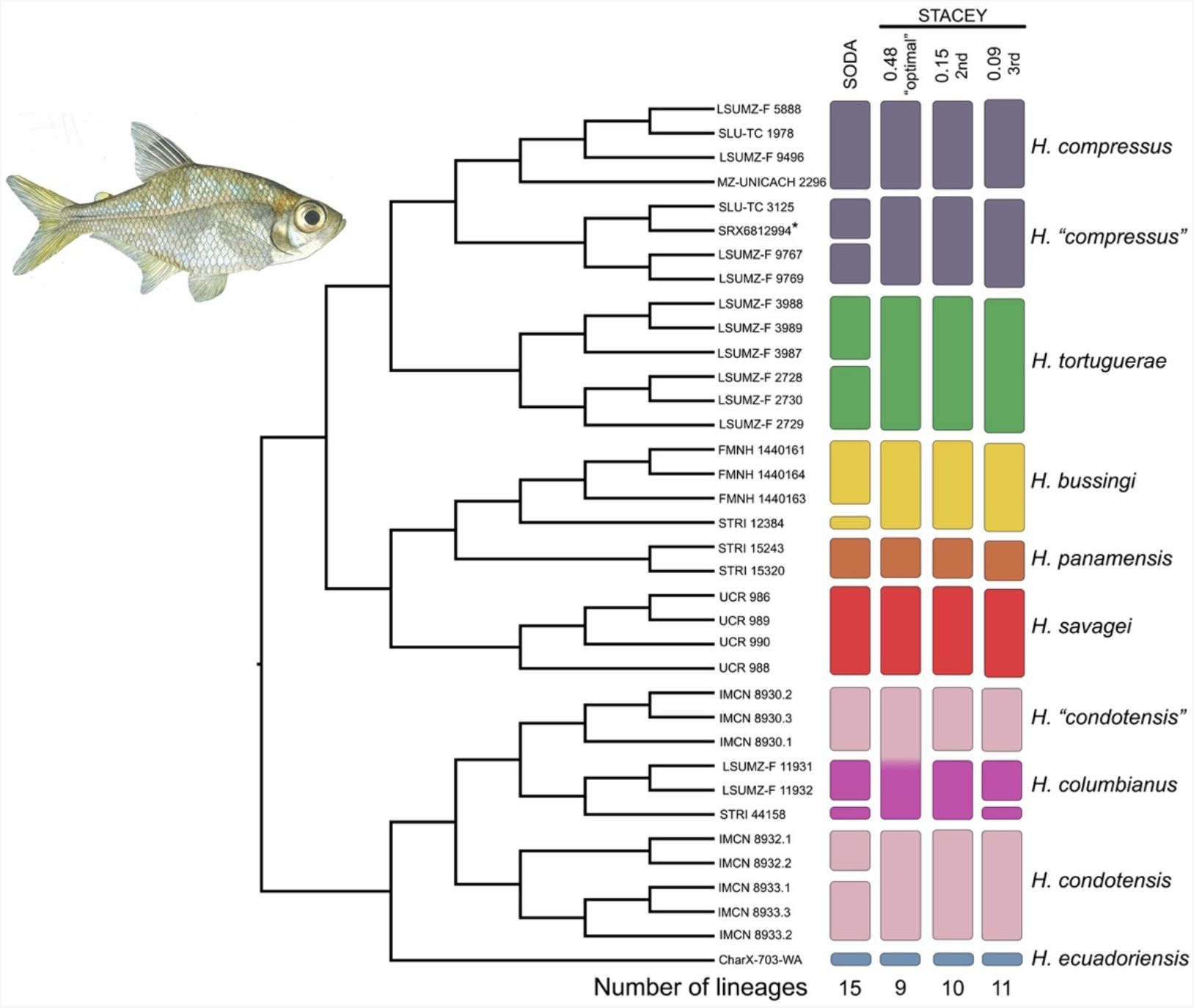
Investigation of cryptic diversity within species of *trans*-Andean *Hyphessobrycon* using a reduced dataset of 36 individuals and no missing data. Cladogram on the left represents the minimum species cluster tree (MSC-Tree) inferred in the STACEY analysis. Tip labels show the catalog number of the samples analyzed in the reduced dataset, asterisks indicate samples from Melo et al. [9] (See S1 Table). The vertical bars to the right represent the output of the species delimitation analyses–number of lineages or clusters and assignment of individuals to each one of them in SODA and STACEY. STACEY 0.48 is the “optimal” number of clusters, STACEY 0.15 is the 2^nd^ alternative number of clusters, and STACEY 0.09 is the 3^rd^ alternative number of clusters recovered. The color scheme of the bars corresponds to the color scheme in figure 1. Illustration of female *Hyphessobrycon compressus* by Duangsamorn Boonwun Boyd.

Both STACEY and SODA congruently uncovered cryptic lineages within samples of *H. compressus* and *H. condotensis* (Fig 4). It is worth noting that the optimal clustering scheme in STACEY (i.e., 0.48) grouped all samples of *H. columbianus* and *H*. “*condotensis*” into a single cluster, in contrast to the two minority schemes (i.e., 0.15 and 0.09) that recovered *H. columbianus* and *H*. “*condotensis*” as independent clusters (Fig 4). For the optimal number of clusters, the probabilities that all samples belong to their assigned “species cluster” were high and ranged from p = 0.79 – 1.0 (Fig 5), with the exception of *H*. “*condotensis*” and *H. columbianus* that showed a probability of p = 0.55 of belonging to the same cluster (Fig 5). The SODA analysis further suggests that there is cryptic diversity within *H*. “*compressus*”, *H. tortuguerae, H. bussingi, H. columbianus*, and *H. condotensis* (Fig 4).

**Figure 5.**
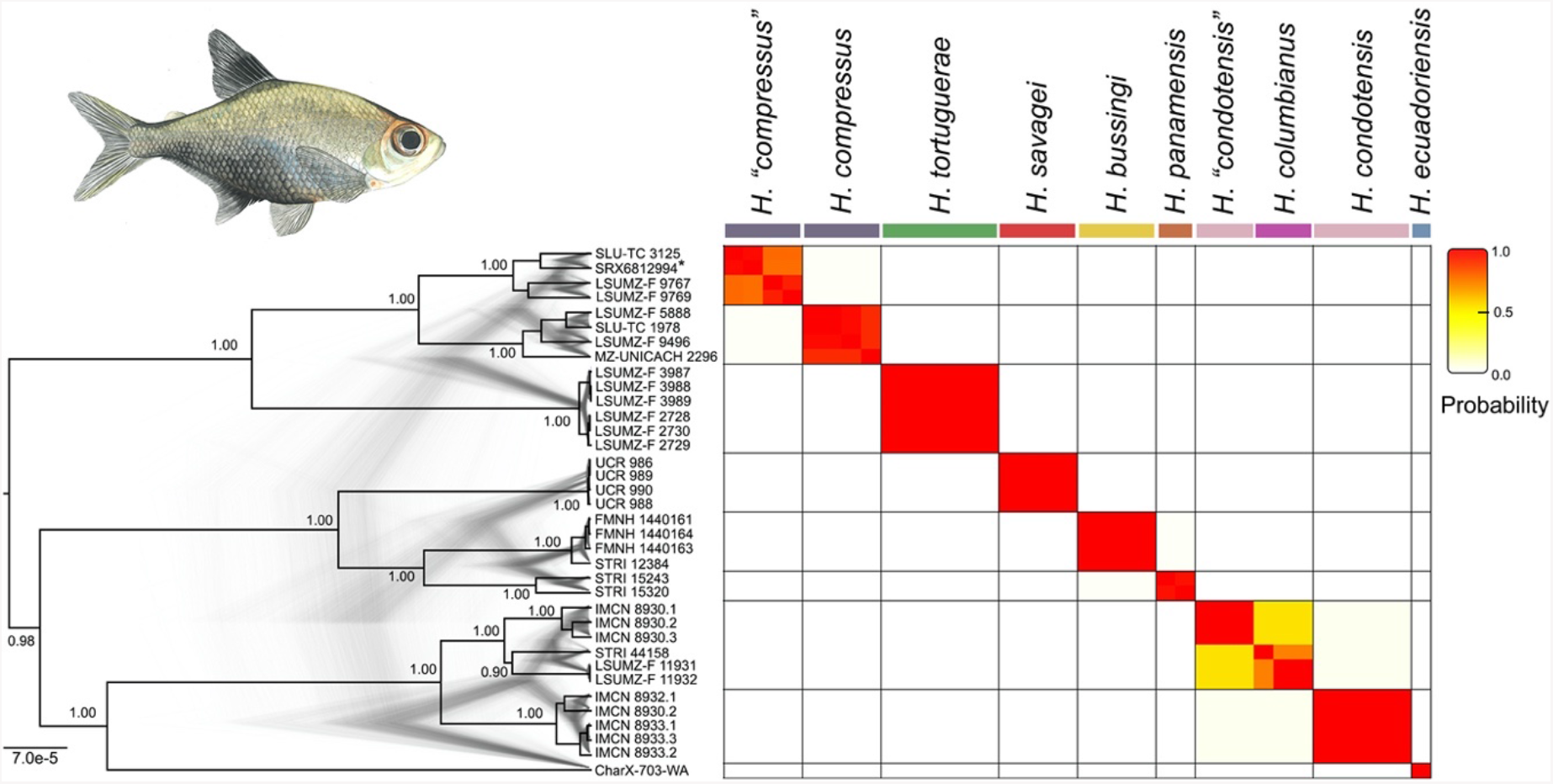
Assignment probability matrix from STACEY delimitation analysis. Each cell in the matrix shows the posterior probability of a pair of samples to belong to the same species cluster. Tip labels show the catalog number of the samples analyzed in the reduced dataset, asterisks indicate sample from Melo et al. [9] (See S1 Table). Darker colors indicate higher probability and lighter colors lower probability. The unrooted minimum species cluster tree (MSC-Tree) of *trans*-Andean *Hyphessobrycon* is shown to the left with posterior probabilities on the nodes. The posterior distribution of trees from the STACEY analysis is superimposed on the MSC-Tree. Illustration of male *Hyphessobrycon compressus* by Duangsamorn Boonwun Boyd.

### Biogeographic Analyses

We identified the DEC-like model as the best fit biogeographical model for our dataset (Table 1). The DEC-like model was the best fit model in both model selection analyses, with and without the +J parameter (Table 1 and S1 Table). Following a recent critique to the usage of the +J parameter (see Ree & Sanmartín [83]) we present our results based on the analysis without the + J parameter (Table 1).

**Table 1.** Parameter estimates of the three biogeographic models tested. LnL = log-likelihood score for the model. *k* = number of parameters, *d* = dispersal rate, *e* = extinction rate or range loss rate AICc = Akaike information criterion corrected. AICc wt = Akaike information criterion corrected weighted. The best biogeographic model was selected based on the highest AICc wt, in bold.

| Model | LnL | $k$ | $d$ | $e$ | AICc | AICc wt |
| --- | --- | --- | --- | --- | --- | --- |
| DEC-like | -25.52 | 2 | 0.57 | 0.43 | 56.24 | **0.62** |
| DIVA-like | -25.99 | 2 | 0.17 | $1.00 \text{ e}^{-12}$ | 57.18 | 0.38 |
| BAYAREA-like | -37.36 | 2 | 0.15 | 0.21 | 79.93 | $4.40 \text{ e}^{-06}$ |

Based on the DEC-like model, we inferred a wide ancestral range east of the Andes and in the Chocó block (AB, Fig 6) for the most recent common ancestor (MRCA) of t*rans*-Andean *Hyphessobrycon* and its sister *cis-*Andean group (node 25 in Fig 6). We inferred six dispersal events and three vicariance events in the clade of *trans*-Andean *Hyphessobrycon*. At crown node 25, we inferred a dispersal event into the Chorotega block and a vicariant event that led to the MRCA of all *trans*-Andean *Hyphessobrycon* (Fig 6). The most likely estimated ancestral range of the MRCA of *trans*-Andean *Hyphessobrycon* included the Chocó and Chorotega blocks (node 22, Fig 6). At node 22, we inferred a vicariant event that lead to the divergence of the Chocó species group and the Central American species of *Hyphessobrycon*. The most likely estimated ancestral ranges of the MRCA of the Chocó species group (node 21) and Central American *Hyphessobrycon* (node 18) were the Chocó and Chorotega blocks, respectively (Fig 6). At node 18, we inferred a dispersal event into the Chortis block and a vicariant event that lead to the split of the ‘panamensis’ species group and the ‘compressus’ species group. The most likely ancestral range of the MRCA of the ‘panamensis’ group (node 17) was the Chorotega, and the Chortis block for the ‘compressus’ species group (node 15) (Fig 6). Subsequently, the MRCA of the ‘compressus’species group dispersed into the Maya block (node 15; Fig 6). Finally, we inferred three dispersal events within the Chorotega block, one at node 16, and two at node 14 (S2 Table).

**Figure 6.**
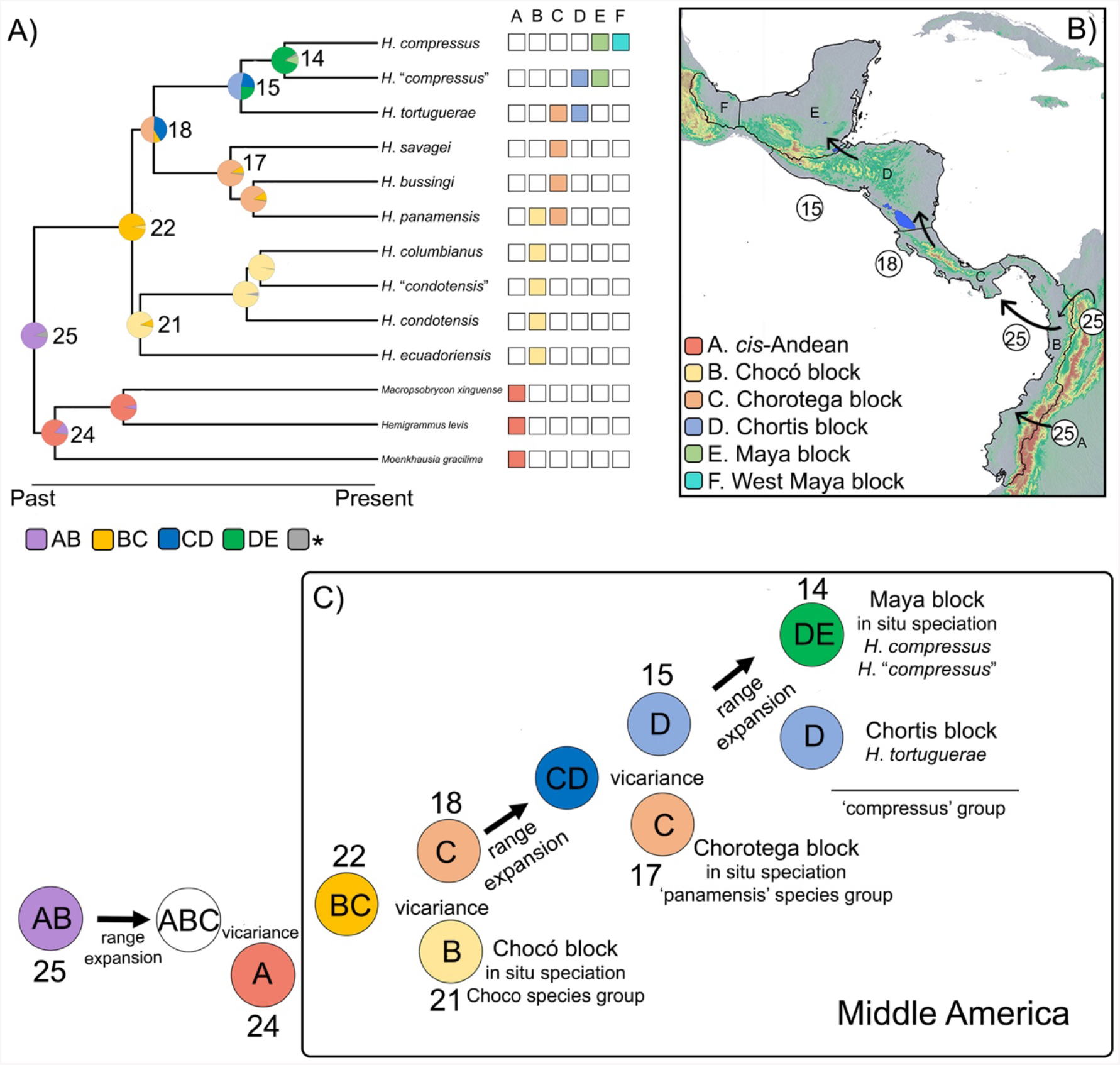
Biogeographic reconstruction of the colonization of Middle America by the genus *Hyphessobrycon* (i.e., *trans*-Andean *Hyphessobrycon*) based on the DEC-like model. A) Ancestral range estimation based on the best biogeographic model DEC-like on the 75% maximum likelihood pruned topology (see methods). Pie charts on nodes represent the probability of the estimated ancestral ranges for each clade. Boxes to the right show present-day presence/absence of *trans*-Andean *Hyphessobrycon* in the geological blocks (i.e., biogeographic units) considered in this analysis. Boxes below the phylogeny show ancestral ranges estimated that encompass two geological blocks, gray with asterisk represent other ancestral ranges. B) Map of Middle America depicting geological blocks and their letter codes. Numbers on circles indicate nodes in A for which we inferred a dispersal event (colonization) to a new geological block. C) Schematic representation of major biogeographic events during the colonization of Middle America by *Hyphessobrycon*. Circles represent nodes on the phylogeny of *trans*-Andean *Hyphessobrycon*, node numbers above or below circles, and their most likely ancestral range estimated (see A), circles without node number represent an inferred event (i.e., range expansion) that occurred in the previous node.

## Discussion

We present the first and most complete phylogenomic hypothesis of *trans*-Andean *Hyphessobrycon* (including *H. compressus*). Our analyses overwhelmingly support the monophyly of *trans*-Andean species of *Hyphessobrycon*, and find a complete lack of evidence supporting the type species of the genus, *H. compressus*, as part of the *cis*-Andean ‘rosy tetra’ clade (Figs 2, 3 and S1 Fig). In agreement with the hypothesis of García-Alzate et al. [25], the phylogenetic placement of *H. compressus*, warrants the *trans*-Andean species to be treated as *Hyphessobrycon sensu stricto*, whereas *cis*-Andean species should be assigned to different genera.

### Phylogenetic relationships

Concatenated and coalescent analyses of 1682 UCE loci are concordant with previous hypotheses that species currently recognized as *Hyphessobrycon* are polyphyletic (Fig 2; e.g., [14, 16, 21]). We recovered strong support for the monophyly of the *trans*-Andean species of *Hyphessobrycon* (Figs 2 - 3 and S1 Fig). A clade comprised of *Macropsobrycon xinguense, Hemigrammus levis*, and *Moenkhausia gracilima* all distributed in the Amazon river basin was inferred as sister to the *trans*-Andean *Hyphessobrycon* clade (Figs 2 and 3). Despite the strong support for the sister relationship of *trans*-Andean *Hyphessobrycon* (LPP = 1, UFBoot2 = 100, SH-aLRT = 100; Figs 2 and S1 Fig), an alternative hypothesis (i.e., *H. diancistrus* + *trans*-Andean *Hyphessobrycon*) was observed in roughly equal frequency (S2 Fig). Recent phylogenomic work of tropical characoids that included a few representatives of *Hyphessobrycon* recovered a clade of *H. compressus* + *H. diancistrus* sister to *M. xinguense, H. levis*, and *M. gracilima* [9]. Furthermore, a close relationship of *H. compressus* to other *cis*-Andean *Hyphessobrycon* species (i.e., *H. bayleyi, H. diancistrus*, and *H. otrynus*) from the Amazon basin has been suggested based on unpublished molecular data, see Lima et al. [84]. Our phylogenomic hypothesis unambiguously placed *H. compressus* in a clade exclusively of *trans*-Andean *Hyphessobrycon* (Figs 2 - 3 and S1 Fig) and challenge previous hypotheses that suggest that *H. compressus* is closely allied with *cis*-Andean taxa [14, 21, 22, 84]. Undoubtedly, future work that includes more taxa of the subfamily Stethaprioninae will enable us to more confidently place the *trans*-Andean *Hyphessobrycon* within the most diverse subfamily of Characidae [20].

### Phylogenetic relationships and cryptic diversity of *trans*-Andean *Hyphessobrycon*

Within the *trans-*Andean *Hyphessobrycon*, our results support the recognition of three subclades: the Chocó species group, the ‘panamensis’ species group, and the ‘compressus’ species group. Our analysis recovered the Chocó species group sister to a clade composed of the ‘panamensis’ + ‘compressus’ species groups (Fig 3A). Furthermore, our species delimitation approaches uncovered cryptic diversity in two species of the Chocó and the ‘compressus’species groups: *H. condotensis* and *H. compressus* respectively (Figs 4 and 5). Furthermore, we interpret the inferred cryptic diversity within *H*. “*compressus”, H. tortuguerae, H. bussingi*, and *H. columbianus* by the SODA analysis (Fig 4) as population-level structure (see methods). The Chocó species group, which is restricted to aquatic systems in the Chocó block in northwestern South America (Fig 1) [61, 62], is composed of *H. ecuadoriensis, H. condotensis, H. columbianus*, and a lineage named *H*. “*condotensis*” (Figs 4 and 5) including individuals from the upper Atrato basin in Colombia. It is worth noting that *H*. “*condotensis*” is closely related to *H. columbianus*, and both lineages are distributed in aquatic systems that drain to the Atlantic slope of northwestern South America. In contrast, samples assigned to *H. condotensis* were collected in the San Juan and Baudo rivers which drain into the Pacific slope of Colombia (Fig 1). Despite the lack of samples of *H. daguae* in our study, we hypothesized that this species belongs to the Chocó species group because of the observed correlation of phylogeny (i.e., species groups) with geographic distributions (i.e., geological blocks). Furthermore, close relationships of *H. daguae* with species of *Hyphessobrycon* in northwestern South America has been proposed based on morphological characters, see Ota et al. [23]. The taxonomy of *Hyphessobrycon* species distributed in the Chocó block has remained in flux [23, 25, 85] and the presence of cryptic diversity has been suggested [86]. Our work uncovered cryptic diversity in the region (Figs 4 and 5) and disagrees with the currently taxonomic hypothesis of four valid species of *Hyphessobrycon* (i.e., *H. columbianus, H. condotensis, H. daguae*, and *H. ecuadoriensis*) distributed in the Chocó block (see Ota et al. [23]). For example, the species *H. sebastiani* was recognized as a junior synonym of *H. condotensis* based on morphological and coloration characters [23]. Our robust phylogenomic hypothesis (Figs 2, 3, and S1 Fig) coupled with the investigation of cryptic diversity (Figs 4 and 5) do not support the recognition of *H. condotensis* as a single widespread species in the Pacific and Atlantic slope of Colombia (Fig 1) proposed by Ota et al. [23]. Our hypothesis is that the cryptic diversity that we uncovered in the upper Atrato basin corresponds to samples of the species *H. sebastiani*, see García-Alzate et al. [25].

Finally, despite the fact that our coalescent and concatenation analyses both recover the monophyly of the Chocó species group (Figs 2 and S1 Fig), gCF and NQS suggest that the placement of *H. ecuadoriensis* is not unambiguously resolved. Specifically, we observed a similar proportion of gene trees that support recognition of the Chocó species group excluding *H. ecuadoriensis*. This hypothesis resolved *H. ecuadoriensis* as the sister to all *trans*-Andean *Hyphessobrycon* (T2; Fig 3D). Future work that includes more robust sampling is needed to better understand and recognize the diversity of *Hyphessobrycon* in the Chocó block and will help to clarify the phylogenetic relationships among this group and within *trans*-Andean *Hyphessobrycon*.

The ‘panamensis’ species group is comprised of three species, *H. savagei, H. panamensis*, and the recently described *H. bussingi* [23]. These three species are mainly distributed in the Pacific (*H. savagei*) and Atlantic (*H. bussingi* and *H. panamensis*) slopes of the Chorotega block [62]. The sister relationship of the two Atlantic species relative to the Pacific one suggests a possible vicariant role of the orogenic process giving rise to the Talamanca mountain range [87] that split the MRCA of the ‘panamensis’ species group into two Pacific and Atlantic lineages. Following divergence, speciation in situ may have occurred within the Atlantic slope of the Chorotega block. The ‘compressus’ species group is comprised of three lineages: *H. compressus, H*. “*compressus*”, and *H. tortuguerae*. The recovered close relationships between these three lineages support the hypothesis of Böhlke [88], who allies *H. compressus, H. tortuguerae*, and *H. milleri* (currently synonymized under *H. compressus*, see below) regardless of differences in coloration among other characters [88]. The uncovering of cryptic diversity within *H. compressus* (Fig 1) is remarkable, considering a recently and thorough systematic work that redescribed *H. compressus* and recognized it as a widespread species (Fig 1), across Mexico, Guatemala and Belize, “without detected variations in meristic and morphometric data” [21]. These results led to the recognition of *H. milleri* as junior synonym of *H. compressus* [21]. Interestingly, our sampling across the distribution of *H. compressus* (Fig 1) mirrors the distribution of specimens analyzed in the previous morphological study (see fig. 7 of Carvalho & Malabarba [21]), which highlights the conserved morphology of some lineages of *trans*-Andean *Hyphessobrycon* (Figs 4 and 5). We propose two plausible hypotheses for the identity of the cryptic lineage *H. “compressus*”: a) it belongs to the previously recognized *H. milleri*, hence extending its distribution into southern and northern Belize, or b) it represents a newly uncovered lineage of the ‘compressus’ species group. Specimens of *Hyphessobrycon* in the Motagua River basin, where the type locality of *H. milleri* is located, have not been collected since 1974 despite recent collecting efforts by the authors (DJE and CDM). The inclusion of samples from the Motagua basin for molecular analyses would be key for testing our proposed hypotheses regarding the identity of the *H*. “*compressus*” lineage.

Finally, the relationships recovered among the three species groups, (Chocó(‘panamensis’,’compressus’)), do not support the recent hypothesis of relationships for some species of *trans*-Andean *Hyphessobrycon* that proposed the ‘*H. panamensis* species-group’ sensu Ota et al. [23] is comprised of *H. bussingi, H. columbianus, H. condotensis, H. daguae, H. panamensis*, and *H. savagei*. Coloration patterns alone or in combination with morphological characters have been used to propose relationships among species of *trans*-Andean *Hyphessobrycon*. For example, it has been suggested that pigmentation (or lack thereof) on the dorsal fin reflects relationships among species of *trans*-Andean *Hyphessobrycon* including the ‘*H. panamensis* species-group’ [21 – 23]. However, our robust phylogenomic hypotheses do not support the monophyly of the ‘*H. panamensis* species-group’ sensu Ota et al. [23] and cast doubt on the use of coloration patterns as phylogenetically informative characters for *trans*-Andean *Hyphessobrycon*. Finally, it is worth noting that for the Central American species of *Hyphessobrycon* (i.e. *H. compressus, H. tortuguerae, H. savagei, H. milleri*, and *H. panamensis*) three morphological synapomorphies that unite these species have been proposed: a) premaxilla with seven teeth in the inner row, b) round foramen in the ventral region of the quadrate, and two foramina in the ventral margin of the epiotic bone, see García-Alzate et al. [25]. Future work is necessary to test if these synapomorphies are diagnostic of *Hyphessobrycon sensu stricto*.

### Colonization of Middle America

The best-fit biogeographical model first inferred a vicariant event separating sister *cis*-Andean taxa and *trans*-Andean *Hyphessobrycon* (node 25; Fig 6). This event was plausibly promoted by orogenic processes in northwestern South America, like the uplift of the Andes during the Miocene [89, 90], that played an important role shaping the riverscape that promoted the origin of *trans*-Andean ichthyofauna [69, 91 – 93].

Our results support a single colonization event of Middle America by *Hyphessobrycon*. The use of geological blocks as biogeographic units allowed us to uncover a stepwise pattern of colonization from South (Chocó block) to North (Maya block), followed by in-situ diversification within each geological block (Fig 6). Our ancestral range estimation inferred that the MRCA of all *trans*-Andean *Hyphessobrycon* was distributed in an area comprising the Chocó and Chorotega blocks (node 22; Fig 6). In this region, another vicariant event led to the separation of the Chocó and ‘panamensis’ species groups. Our biogeographical inferences together with the present-day distribution of the Chocó and ‘panamensis’ species groups agree with the early colonization model of ostariophysan fishes of Bermingham and Martin [94]. This model suggests an early colonization into Central America (Chorotega block) during the late Miocene, followed by extinction (in some taxa) in some river basins due to changes in sea levels during the Pliocene [68, 69, 91, 93 – 96]. Furthermore, our current understanding of the complex landscape in lower Middle America and the timing of the initial closure of the Isthmus of Panama (∼ 20 million years ago) [67, 97] provides evidence of connectivity between the Chocó and Chorotega blocks during the Miocene.

From the Chorotega block, we inferred a range expansion into the Chortis block followed by a vicariant event that separated the ‘panamensis’ and ‘compressus’ species groups. The range fragmentation of the MRCA of the Central American species of *Hyphessobrycon* may be explained by a marine corridor across the Nicaragua depression, which was located between the Chorotega and Chortis blocks until the Pliocene [63]. Global sea level changes [98] have impacted the landscape of Middle America (see Bagley and Johnson [99] and references therein). Due to physiological constraints of *Hyphessobrycon* (i.e., salinity intolerance), a marine corridor likely extirpated ancestral populations and limited gene flow between populations on either side of the barrier that gave origin to the ‘panamensis’ and ‘compressus’ groups. Finally, present-day disjunct distributions between species in the ‘compressus’ group (Fig 1) preclude us from making inferences regarding the colonization route from the Chortis into the Maya block.

## Conclusions

Historically, it has been challenging to resolve the phylogenetic relationships in some lineages of the highly diverse Characidae, in part due to their conserved morphologies [14, 22, 100] or homoplasy of phylogenetic characters [22, 69, 101]. Our robust phylogenomic hypothesis resolves the placement of *H. compressus* together with the other trans-Andean species, instead of with the *cis*-Andean species of the ‘rosy tetra’ clade where it had been historically grouped [21, 22]. Our hypothesis supports that *Hyphessobrycon* sensu stricto should be restricted to *trans*-Andean taxa. Future work that incorporates more taxa of Stethaprioninae will help us to more confidently place *Hyphessobrycon* sensu stricto within this subfamily.

Our results bring into question the utility of coloration patterns as phylogenetically informative characters for *trans*-Andean *Hyphessobrycon*. The unveiling of cryptic diversity highlights the conserved morphology among divergent lineages of *trans*-Andean *Hyphessobrycon*. Finally, we propose an interesting biogeographic model for the colonization of Middle America by ostariophysan fishes, in which the ancestral lineages first colonized the geological blocks in a stepwise fashion, and after colonization, in-situ speciation took place within each geological block.

## Acknowledgments

We would like to thank Consejo Nacional of Areas Protegidas (CONAP) for granting research and collecting permits to DJE (DRM/002/2018; 48-2018), especially to Airam López, Héisel Arriola, and MSc José Luis Echeverría for their instrumental assistance through this process. CDM thanks the Negaunee Foundation and Grainger Bioinformatics Center at the Field Museum for their support of his work. DJE thanks César Fuentes, Lucia López, Marvin Xajil, Elyse Parker, Ricardo Penagos, Francis Santos for their assistance in the field. We thank Dr. Wilfredo Matamoros (UNICACH), Dr. Jonathan Armbruster (Auburn University), Dr. Arturo Angulo (Universidad de Costa Rica), Dr. Windsor Aguirre (De Paul University), Smithsonian Tropical Institute, Dr. Kyle Piller (Southeastern Louisiana University), Dr. Luke Bower (Clemson University) for facilitate samples used in this work. We thank Dr. Jeremy Brown (Louisiana State University) and Elyse Parker (Yale University) for their valuable comments on this work. DJE thanks Duangsamorn Boonwun Boyd for their watercolors of *Hyphessobrycon compressus* used in this work.

## Data Availability Statement

The generated raw reads are archived in the sequence repository archive (SRA) of the National Center for Biotechnology Information (NCBI) BioProject XXXXXXXXXXX

## Ethics Statement

Research and collecting permits granted to DJE (DRM/002/2018; 48-2018) by Consejo Nacional de Areas Protegidas de Guatemala (CONAP).

## References

1. Albert JS, Reis RE. Introduction to Neotropical Freshwaters. Historical Biogeography of Neotropical Freshwater Fishes. 2011. pp. 2–19. doi:10.1525/california/9780520268685.003.0001

2. Albert JS, Tagliacollo VA, Dagosta F. Diversification of Neotropical Freshwater Fishes. Annu Rev Ecol Evol Syst. 2020. pp. 27–53. doi:10.1146/annurev-ecolsys-011620-031032

3. Chakrabarty P, Faircloth BC, Alda F, Ludt WB, Mcmahan CD, Near TJ, et al. Phylogenomic Systematics of Ostariophysan Fishes: Ultraconserved Elements Support the Surprising Non-Monophyly of Characiformes. Syst Biol. 2017;66: 881–895.

4. Arcila D, Ortí G, Vari R, Armbruster JW, Stiassny MLJ, Ko KD, et al. Genome-wide interrogation advances resolution of recalcitrant groups in the tree of life. Nat Ecol Evol. 2017;1: 20.

5. Ilves KL, Torti D, López-Fernández H. Exon-based phylogenomics strengthens the phylogeny of Neotropical cichlids and identifies remaining conflicting clades (Cichliformes: Cichlidae: Cichlinae). Mol Phylogenet Evol. 2018;118: 232–243.

6. Alda F, Tagliacollo VA, Bernt MJ, Waltz BT, Ludt WB, Faircloth BC, et al. Resolving Deep Nodes in an Ancient Radiation of Neotropical Fishes in the Presence of Conflicting Signals from Incomplete Lineage Sorting. Syst Biol. 2019. pp. 573–593. doi:10.1093/sysbio/syy085

7. Alda F, Ludt WB, Elías DJ, McMahan CD, Chakrabarty P. Comparing Ultraconserved Elements and Exons for Phylogenomic Analyses of Middle American Cichlids: When Data Agree to Disagree. Genome Biol Evol. 2021;13. doi:10.1093/gbe/evab161

8. Ochoa LE, Datovo A, DoNascimiento C, Roxo FF, Sabaj MH, Chang J, et al. Phylogenomic analysis of trichomycterid catfishes (Teleostei: Siluriformes) inferred from ultraconserved elements. Sci Rep. 2020;10: 2697.

9. Melo BF, Sidlauskas BL, Near TJ, Roxo FF, Ghezelayagh A, Ochoa LE, et al. Accelerated Diversification Explains the Exceptional Species Richness of Tropical Characoid Fishes. Syst Biol. 2021;71: 78–92.

10. Silva GSC, Roxo FF, Melo BF, Ochoa LE, Bockmann FA, Sabaj MH, et al. Evolutionary history of Heptapteridae catfishes using ultraconserved elements (Teleostei, Siluriformes). Zool Scr. 2021. pp. 543–554. doi:10.1111/zsc.12493

11. Souza CS, Melo BF, Mattox GMT, Oliveira C. Phylogenomic analysis of the Neotropical fish subfamily Characinae using ultraconserved elements (Teleostei: Characidae). Mol Phylogenet Evol. 2022;171: 107462.

12. Oliveira C, Avelino GS, Abe KT, Mariguela TC, Benine RC, Ortí G, et al. Phylogenetic relationships within the speciose family Characidae (Teleostei: Ostariophysi: Characiformes) based on multilocus analysis and extensive ingroup sampling. BMC Evol Biol. 2011;11: 275.

13. Mirande JM. Weighted parsimony phylogeny of the family Characidae (Teleostei: Characiformes). Cladistics. 2009;25: 574–613.

14. Mirande JM. Morphology, molecules and the phylogeny of Characidae (Teleostei, Characiformes). Cladistics. 2019;35: 282–300.

15. Dagosta FCP, De Pinna M. The Fishes of the Amazon: Distribution and Biogeographical Patterns, with a Comprehensive List of Species. Bull Am Mus Nat Hist. 2019. p. 1. doi:10.1206/0003-0090.431.1.1

16. Guimarães EC, de Brito PS, Feitosa LM, Costa LFC, Ottoni FP. A new cryptic species of Hyphessobrycon Durbin, 1908 (Characiformes, Characidae) from the Eastern Amazon, revealed by integrative taxonomy. Zoosyst Evol. 2019. pp. 345–360. doi:10.3897/zse.95.34069

17. García-Alzate CA, Lima F, Taphorn DC, Mojica JI, Urbano-Bonilla A, Teixeira TF. A new species of Hyphessobrycon Durbin (Characiformes: Characidae) from the western Amazon basin in Colombia and Peru. J Fish Biol. 2020;96: 1444–1453.

18. Faria TC, Guimarães KLA, Rodrigues LRR, Oliveira C, Lima FCT. A new Hyphessobrycon (Characiformes: Characidae) of the Hyphessobrycon heterorhabdus species-group from the lower Amazon basin, Brazil. Neotrop Ichthyol. 2021. doi:10.1590/1982-0224-2020-0102

19. Dagosta FCP, Seren TJ, Ferreira A, Marinho MMF. The emerald green tetra: a new restricted-range Hyphessobrycon (Characiformes: Characidae) from the upper rio Juruena, Chapada dos Parecis, Brazil. Neotrop Ichthyol. 2022. doi:10.1590/1982-0224-2021-0119

20. Fricke R, Eschmeyer WN, and van der Laan R. (eds). Eschmeyer’s catalog of fishes: genera, species, references. 2022. Electronic version accessed 20/ February/ 2022 http://researcharchive.calacademy.org/research/ichthyology/catalog/fishcatmain.asp

21. Carvalho FR, Malabarba LR. Redescription and osteology of Hyphessobrycon compressus (Meek) (Teleostei: Characidae), type species of the genus. Neotrop Ichthyol. 2015. pp. 513–540. doi:10.1590/1982-0224-20140173

22. Weitzman SH, and Palmer L. A new species of Hyphessobrycon (Teleostei: Characidae) from the Neblina region of Venezuela and Brazil, with comments on the putative ‘rosy tetra clade.’ Ichthyol Explor Freshw. 1997; 7, 209–242.

23. Ota RR, Carvalho FR, Pavanelli CS. Taxonomic review of the Hyphessobrycon panamensis species-group (Characiformes: Characidae). Zootaxa. 2020;4751: zootaxa.4751.3.1.

24. Meek SE. The freshwater fishes of Mexico north of the isthmus of Tehuantepec. Field Columbian Museum, Zoölogical series. 1904, v. 5: i–lxiii.

25. García Alzate CA, Román-Valencia C, Taphorn DC. Una nueva especie de Hyphessobrycon (Characiformes: Characidae) de la cuenca del río Telembí, vertiente sur del Pacífico, Colombia. Rev Biol Trop. 2013. doi:10.15517/rbt.v61i1.10944

26. Faircloth BC, McCormack JE, Crawford NG, Harvey MG, Brumfield RT, Glenn TC. Ultraconserved elements anchor thousands of genetic markers spanning multiple evolutionary timescales. Syst Biol. 2012;61: 717–726.

27. Sabaj MH. Codes for Natural History Collections in Ichthyology and Herpetology. Copeia. 2020. doi:10.1643/asihcodons2020

28. Ward RD, Hanner R, Hebert PDN. The campaign to DNA barcode all fishes, FISH-BOL. J Fish Biol. 2009;74: 329–356.

29. Altschul SF, Gish W, Miller W, Myers EW, Lipman DJ. Basic local alignment search tool. J Mol Biol. 1990. pp. 403–410. doi:10.1016/s0022-2836(05)80360-2

30. Faircloth BC, Glenn TC. Not all sequence tags are created equal: designing and validating sequence identification tags robust to indels. PLoS One. 2012;7: e42543.

31. Burress ED, Alda F, Duarte A, Loureiro M, Armbruster JW, Chakrabarty P. Phylogenomics of pike cichlids (Cichlidae: Crenicichla): the rapid ecological speciation of an incipient species flock. J Evol Biol. 2018;31: 14–30.

32. Faircloth BC, Alda F, Hoekzema K, Burns MD, Oliveira C, Albert JS, et al. A Target Enrichment Bait Set for Studying Relationships among Ostariophysan Fishes. Copeia. 2020. p. 47. doi:10.1643/cg-18-139

33. Faircloth BC. PHYLUCE is a software package for the analysis of conserved genomic loci. Bioinformatics. 2016;32: 786–788.

34. Bolger AM, Lohse M, Usadel B. Trimmomatic: a flexible trimmer for Illumina sequence data. Bioinformatics. 2014;30: 2114–2120.

35. Chen S, Zhou Y, Chen Y, Gu J. fastp: an ultra-fast all-in-one FASTQ preprocessor. Bioinformatics. 2018. pp. i884–i890. doi:10.1093/bioinformatics/bty560

36. Handika H. YAP: A single executable pipeline for phylogenomics (v0.3.0). Zenodo. 2022. https://doi.org/10.5281/zenodo.6128874

37. Bankevich A, Nurk S, Antipov D, Gurevich AA, Dvorkin M, Kulikov AS, et al. SPAdes: a new genome assembly algorithm and its applications to single-cell sequencing. J Comput Biol. 2012;19: 455–477.

38. Prjibelski A, Antipov D, Meleshko D, Lapidus A, Korobeynikov A. Using SPAdes De Novo Assembler. Curr Protoc Bioinformatics. 2020;70: e102.

39. Katoh K, Standley DM. MAFFT multiple sequence alignment software version 7: improvements in performance and usability. Mol Biol Evol. 2013;30: 772–780.

40. Castresana J. Selection of conserved blocks from multiple alignments for their use in phylogenetic analysis. Mol Biol Evol. 2000;17: 540–552.

41. Handika H, Esselstyn J. SEGUL: An ultrafast, memory-efficient alignment manipulation and summary tool for phylogenomics. doi:10.22541/au.165167823.30911834/v1

42. Minh BQ, Schmidt HA, Chernomor O, Schrempf D, Woodhams MD, von Haeseler A, et al. IQ-TREE 2: New Models and Efficient Methods for Phylogenetic Inference in the Genomic Era. Mol Biol Evol. 2020;37: 1530–1534.

43. Kalyaanamoorthy S, Minh BQ, Wong TKF, von Haeseler A, Jermiin LS. ModelFinder: fast model selection for accurate phylogenetic estimates. Nat Methods. 2017;14: 587–589.

44. Hoang DT, Chernomor O, von Haeseler A, Minh BQ, Vinh LS. UFBoot2: Improving the Ultrafast Bootstrap Approximation. Mol Biol Evol. 2018;35: 518–522.

45. Guindon S, Dufayard J-F, Lefort V, Anisimova M, Hordijk W, Gascuel O. New Algorithms and Methods to Estimate Maximum-Likelihood Phylogenies: Assessing the Performance of PhyML 3.0. Syst Biol. 2010. pp. 307–321. doi:10.1093/sysbio/syq010

46. Zhang C, Rabiee M, Sayyari E, Mirarab S. ASTRAL-III: polynomial time species tree reconstruction from partially resolved gene trees. BMC Bioinformatics. 2018;19: 153.

47. Simmons MP, Gatesy J. Collapsing dubiously resolved gene-tree branches in phylogenomic coalescent analyses. Mol Phylogenet Evol. 2021;158: 107092.

48. Junier T, Zdobnov EM. The Newick utilities: high-throughput phylogenetic tree processing in the UNIX shell. Bioinformatics. 2010;26: 1669–1670.

49. Brown JM, Thomson RC. Bayes Factors Unmask Highly Variable Information Content, Bias, and Extreme Influence in Phylogenomic Analyses. Syst Biol. 2017;66: 517–530.

50. Thomson RC, Brown JM. On the Need for New Measures of Phylogenomic Support. Syst Biol. 2022;71: 917–920.

51. Baum DA. Concordance trees, concordance factors, and the exploration of reticulate genealogy. Taxon. 2007. pp. 417–426. doi:10.1002/tax.562013

52. Sayyari E, Mirarab S. Fast Coalescent-Based Computation of Local Branch Support from Quartet Frequencies. Mol Biol Evol. 2016;33: 1654–1668.

53. Jones G, Aydin Z, Oxelman B. DISSECT: an assignment-free Bayesian discovery method for species delimitation under the multispecies coalescent. Bioinformatics. 2015;31: 991–998.

54. Jones G. Algorithmic improvements to species delimitation and phylogeny estimation under the multispecies coalescent. J Math Biol. 2017;74: 447–467.

55. Rabiee M, Mirarab S. SODA: Multi-locus species delimitation using quartet frequencies. Bioinformatics. 2021. doi:10.1093/bioinformatics/btaa1010

56. Drummond AJ, Rambaut A. BEAST: Bayesian evolutionary analysis by sampling trees. BMC Evol Biol. 2007. p. 214. doi:10.1186/1471-2148-7-214

57. Drummond AJ, Suchard MA, Xie D, Rambaut A. Bayesian phylogenetics with BEAUti and the BEAST 1.7. Mol Biol Evol. 2012;29: 1969–1973.

58. R Core Team. R: A language and environment for statistical computing. 2019. Vienna, Austria, R Foundation for Statistical Computing

59. Winker K. Middle America, not Mesoamerica, is the Accurate Term for Biogeography. Condor. 2011. pp. 5–6. doi:10.1525/cond.2011.100093

60. Gutiérrez-García TA, Vázquez-Domínguez E. Consensus between genes and stones in the biogeographic and evolutionary history of Central America. Quat Res. 2013. pp. 311–324. doi:10.1016/j.yqres.2012.12.007

61. Dengo G. Mid America: Tectonic Setting for the Pacific Margin from Southern Mexico to Northwestern Colombia. The Ocean Basins and Margins. 1985. pp. 123–180. doi:10.1007/978-1-4613-2351-8_4

62. Marshall J. Geomorphology and physiographic provinces. Central America. 2007. doi:10.1201/9780203947043.pt2

63. Coates A, and Obando J. The geologic evolution of the Central American Isthmus. In J.B.C. Jackson, A.F. Budd, & A.G. Coates (Eds), Evolution and environment in tropical America. 1996. pp 21–56. Chicago, The University of Chicago Press

64. Iturralde-Vinent MA, and MacPhee RD. Paleogeography of the Caribbean region: implications for Cenozoic biogeography. Bull Am Mus Nat Hist. 1999; 238

65. Iturralde-Vinent MA. Meso-Cenozoic Caribbean Paleogeography: Implications for the Historical Biogeography of the Region. Intl Geol Rev. 2006. pp. 791–827. doi:10.2747/0020-6814.48.9.791

66. Myers GS. Derivation of the Freshwater Fish Fauna of Central America. Copeia. 1966. p. 766. doi:10.2307/1441405

67. Bacon CD, Silvestro D, Jaramillo C, Smith BT, Chakrabarty P, Antonelli A. Biological evidence supports an early and complex emergence of the Isthmus of Panama. Proc Natl Acad Sci U S A. 2015;112: 6110–6115.

68. Reeves RG, Guy Reeves R, Bermingham E. Colonization, population expansion, and lineage turnover: phylogeography of Mesoamerican characiform fish. Biol J Linn Soc Lond. 2006. pp. 235–255. doi:10.1111/j.1095-8312.2006.00619.x

69. Ornelas-García CP, Domínguez-Domínguez O, Doadrio I. Evolutionary history of the fish genus Astyanax Baird & Girard (1854) (Actinopterygii, Characidae) in Mesoamerica reveals multiple morphological homoplasies. BMC Evol Biol. 2008;8: 340.

70. GBIF.org (21 February 2022) GBIF Occurrence Available from: https://doi.org/10.15468/dl.ch24m5

71. QGIS Geographic Information System. Open Source Geospatial Foundation Project. URL https://qgis.org

72. Lehner B, Grill G. Global river hydrography and network routing: baseline data and new approaches to study the world’s large river systems. Hydrol Process. 2013. pp. 2171–2186. doi:10.1002/hyp.9740

73. Linke S, Lehner B, Ouellet Dallaire C, Ariwi J, Grill G, Anand M, et al. Global hydro-environmental sub-basin and river reach characteristics at high spatial resolution. Sci Data. 2019;6: 283.

74. Paradis E, Claude J, Strimmer K. APE: Analyses of Phylogenetics and Evolution in R language. Bioinformatics. 2004. pp. 289–290. doi:10.1093/bioinformatics/btg412

75. Popescu A-A, Huber KT, Paradis E. ape 3.0: New tools for distance-based phylogenetics and evolutionary analysis in R. Bioinformatics. 2012. pp. 1536–1537. doi:10.1093/bioinformatics/bts184

76. Matzke NJ. Model selection in historical biogeography reveals that founder-event speciation is a crucial process in Island Clades. Syst Biol. 2014;63: 951–970.

77. Yu Y, Blair C, He X. RASP 4: Ancestral State Reconstruction Tool for Multiple Genes and Characters. Mol Biol Evol. 2020;37: 604–606.

78. Bishop P. Drainage rearrangement by river capture, beheading and diversion. Prog Phys Geogr: Earth Environ. 1995. pp. 449–473. doi:10.1177/030913339501900402

79. Dias MS, Oberdorff T, Hugueny B, Leprieur F, Jézéquel C, Cornu J-F, et al. Global imprint of historical connectivity on freshwater fish biodiversity. Ecol Lett. 2014;17: 1130–1140.

80. Ronquist F. Dispersal-Vicariance Analysis: A New Approach to the Quantification of Historical Biogeography. Syst Biol. 1997. pp. 195–203. doi:10.1093/sysbio/46.1.195

81. Ree RH, Smith SA. Maximum likelihood inference of geographic range evolution by dispersal, local extinction, and cladogenesis. Syst Biol. 2008;57: 4–14.

82. Landis MJ, Matzke NJ, Moore BR, Huelsenbeck JP. Bayesian analysis of biogeography when the number of areas is large. Syst Biol. 2013;62: 789–804.

83. Ree RH, Sanmartín I. Conceptual and statistical problems with the DEC J model of founder-event speciation and its comparison with DEC via model selection. J Biogeogr. 2018. pp. 741–749. doi:10.1111/jbi.13173

84. Lima FCT, Bastos DA, Py-Daniel LHR, Ota RP. A new sexually dimorphic Hyphessobrycon from the western Amazon basin (Characiformes: Characidae). Zootaxa. 2022;5116: 253–266.

85. García–Alzate CA, Román–Valencia C, Taphorn DC. A new species of Hyphessobrycon (Teleostei: Characiformes: Characidae) from the San Juan River drainage, Pacific versant of Colombia. Zootaxa. 2010. p. 55. doi:10.11646/zootaxa.2349.1.4

86. Jiménez-Prado P, Aguirre W, Laaz-Moncayo E, Navarrete-Amaya R, Nugra-Salazar F, Rebolledo-Monsalve E, et al. Guía de peces para aguas continentales en la vertienete occidental del Ecuador. Pontificia Universidad Católica del Ecuador Sede Esmeraldas (PUCESE); Universidad del Azuay (UDA) y Museo Ecuatoriano de Ciencias Naturales (MECN) del Instituto Nacional de Biodiversidad. Esmeraldas, Ecuador. 2015. 416 pp.

87. de Boer JZ, Drummond MS, Bordelon MJ, Defant MJ, Bellon H, et al. Cenozoic magmatic phases of the Costa Rican island arc (Cordillera de Talamanca). Geol Soc Am Special Papers. 1995. pp. 35–56. doi:10.1130/spe295-p35

88. Böhlke JE. Studies on Fishes of the family Characidae No. 16. -A New Hyphessobrycon from Costa Rica. Bulletin of the Florida State Museum, Biological Sciences. 1958; v. 3 (no. 4).

89. Hoorn C, Guerrero J, Sarmiento GA, Lorente MA. Andean tectonics as a cause for changing drainage patterns in Miocene northern South America. Geology. 1995. p. 237. doi:10.1130/0091-7613(1995)023<0237:ataacf>2.3.co;2

90. Hoorn C, Wesselingh FP, Hovikoski J, Guerrero J. The Development of the Amazonian Mega-Wetland (Miocene; Brazil, Colombia, Peru, Bolivia). Amazonia: Landscape and Species Evolution. 2011. pp. 123–142. doi:10.1002/9781444306408.ch8

91. Perdices A, Bermingham E, Montilla A, Doadrio I. Evolutionary history of the genus Rhamdia (Teleostei: Pimelodidae) in Central America. Mol Phylogenet Evol. 2002;25: 172–189.

92. Albert JS, Lovejoy NR, and Crampton WGR. Miocene tectonism and the separation of cis- and trans-Andean river basins: Evidence from Neotropical fishes. J South Am Earth Sci. 2006; (Vol. 21, Issues 1-2, pp. 14–27). https://doi.org/10.1016/j.jsames.2005.07.010

93. Picq S, Alda F, Krahe R, Bermingham E. Miocene and Pliocene colonization of the Central American Isthmus by the weakly electric fish Brachyhypopomus occidentalis (Hypopomidae, Gymnotiformes). J Biogeogr. 2014. pp. 1520–1532. doi:10.1111/jbi.12309

94. Bermingham E, Martin AP. Comparative mtDNA phylogeography of neotropical freshwater fishes: testing shared history to infer the evolutionary landscape of lower Central America. Mol Ecol. 1998;7: 499–517.

95. Lovejoy NR, Lester K, Crampton WGR, Marques FPL, Albert JS. Phylogeny, biogeography, and electric signal evolution of Neotropical knifefishes of the genus Gymnotus (Osteichthyes: Gymnotidae). Mol Phylogenet Evol. 2010;54: 278–290.

96. Smith SA, Bermingham E. The biogeography of lower Mesoamerican freshwater fishes. J Biogeogr. 2005. pp. 1835–1854. doi:10.1111/j.1365-2699.2005.01317.x

97. Montes C, Cardona A, McFadden R, Moron SE, Silva CA, Restrepo-Moreno S, et al. Evidence for middle Eocene and younger land emergence in central Panama: Implications for Isthmus closure. Geol Soc Am Bull. 2012. pp. 780–799. doi:10.1130/b30528.1

98. Haq BU, Hardenbol J, Vail PR. Chronology of fluctuating sea levels since the triassic. Science. 1987;235: 1156–1167.

99. Bagley JC, Johnson JB. Phylogeography and biogeography of the lower Central American Neotropics: diversification between two continents and between two seas. Biol Rev Camb Philos Soc. 2014;89: 767–790.

100. Terán GE, Benitez MF, Marcos Mirande J. Opening the Trojan horse: phylogeny of Astyanax, two new genera and resurrection of Psalidodon (Teleostei: Characidae). Zool J Linn Soc. 2020. doi:10.1093/zoolinnean/zlaa019

101. Mirande JM. Phylogeny of the family Characidae (Teleostei: Characiformes): from characters to taxonomy. Neotrop Ichthyol. 2010. pp. 385–568. doi:10.1590/s1679-62252010000300001

