## Supplementary material for "Phylogenomics of *trans*-Andean tetras of the genus *Hyphessobrycon* Durbin 1908 and colonization patterns of Middle America": S2 Table, S1 Figure, and S2 Figure

**S1 Table List of all samples included in this study and summary statistics of the sequencing output recovered for each one of them**. Samples in bold were used for the cryptic diversity analyses. Museum codes follow Sabaj (2020). Excel File

**S2 Table Parameters estimates of six different biogeographic models.** LnL = log-likelihood score for the model. *k* = number of parameters, *d* = dispersal rate, *e* = extinction rate or range loss rate, *j* = jum-dispersal rate, AICc = Akaike information criterion corrected. AICc wt = Akaike information criterion corrected weighted. The best biogeographic model was selected based on the highest AICc wt, in bold.

| Model | LnL | *k* | *d* | *e* | *j* | AICc | AICc wt |
| --- | --- | --- | --- | --- | --- | --- | --- |
| **DEC-like** | -25.52 | 2 | 0.57 | 0.43 | 0 | **56.24** | **0.36** |
| DEC-like + J | -24.58 | 3 | 0.091 | 1.00 e^-12^ | 0.031 | 57.83 | 0.16 |
| DIVA-like | -25.99 | 2 | 0.17 | 1.00 e^-12^ | 0 | 57.18 | 0.22 |
| DIVA-like + J | -24.19 | 3 | 0.10 | 1.00 e^-12^ | 0.028 | 57.05 | 0.24 |
| BAYAREA-like | -37.36 | 2 | 0.15 | 0.21 | 0 | 79.93 | 2.60 e^-06^ |
| BAYAREA-like + J | -27.11 | 3 | 0.079 | 1.00 e^-07^ | 0.054 | 62.89 | 0.013 |

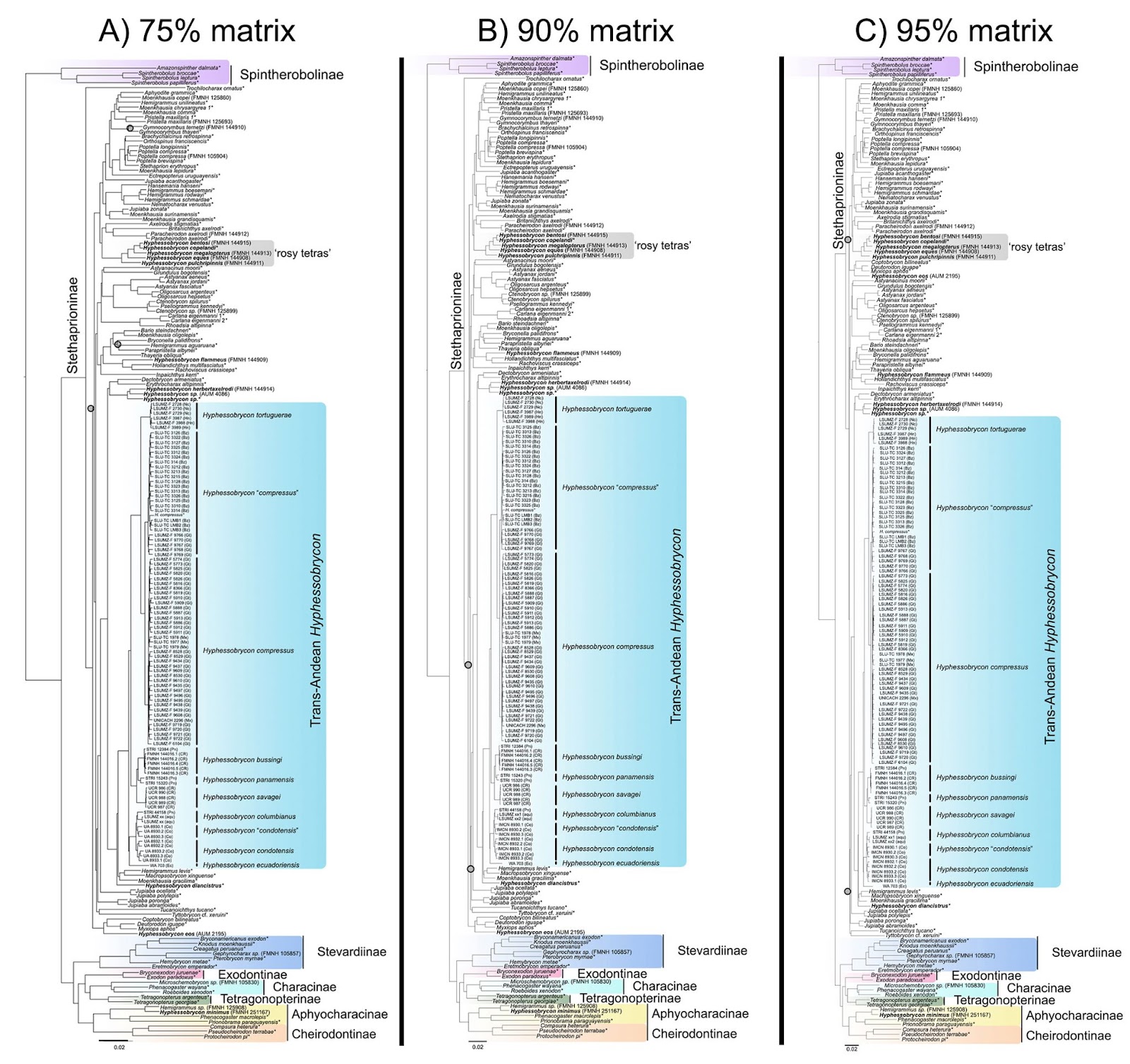

**S1 Figure. Phylogenomic relationships of *trans*-Andean *Hyphessobrycon* based on concatenated analysis of ultraconserved elements**. A) Inferred phylogeny based on the 75% complete data matrix B) inferred phylogeny based on the 90% complete data matrix, and C) inferred phylogeny based on the 75% complete data matrix. All nodes are supported with ultrafast bootstrap (UFBoot2) = 100 and SH-like approximate ratio test (SH-aLRT) = 100 unless noted. Nodes with gray circles UFBoot2 < 90 and SH-aLRT < 90. Species names with asterisk indicates samples from Melo et al. [9]. Species of *Hyphessobrycon* has been bolded.

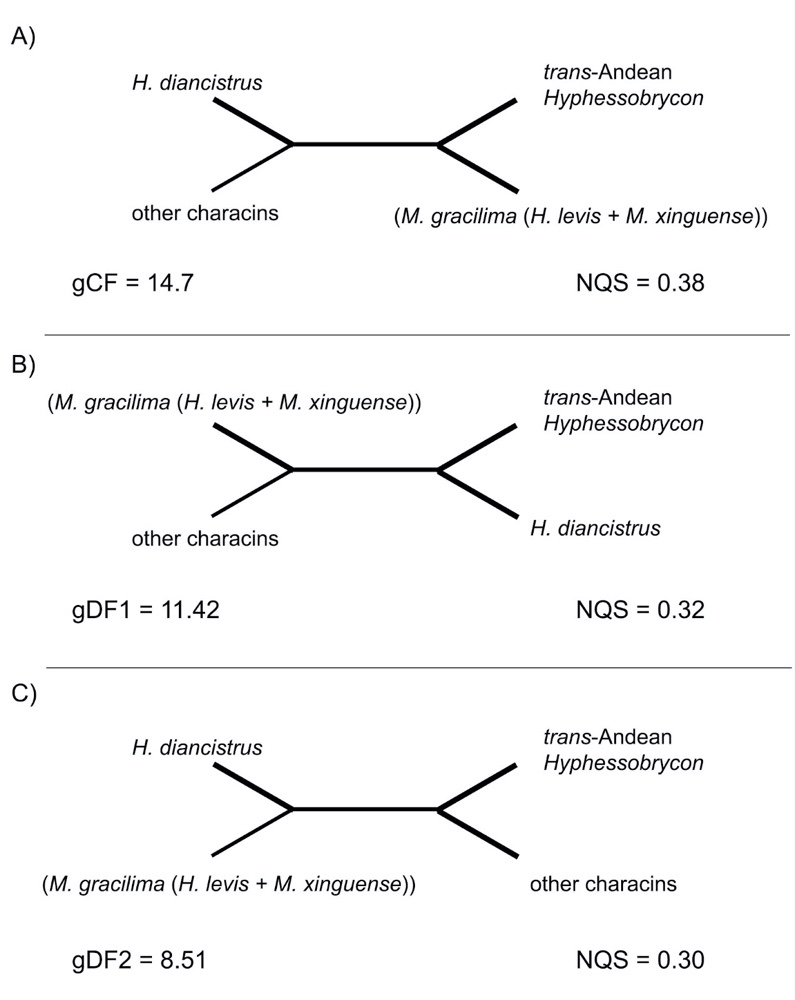

**S2 Figure**. Gene concordance factors (gCF) and normalized quartet scores (NQS) for the main topology/quartet (A) and the two alternatives quartets (B & C) for the inferred sister relationships of *trans*-Andean *Hyphessobrycon*. A) *trans*-Andean *Hyphessobrycon* sister to a clade comprise of *M*. *gracilima*, *H*. *levis*, and *M*. *xinguense* (see Fig. 2) B) *trans*-Andean *Hyphessobrycon* sister to *H*. *diancistrus*, and C) *trans*-Andean *Hyphessobrycon* sister other characins of the subfamily Stethaprioninae. Number of trees decisive for the branch (gN) = 1646, effective number of genes for a branch of interest (EN) = 1220.72. gCF = gene concordance factor, gDF1 = gene discordance factor for NNI-1 branch, gDF2 = gene discordance factor for NNI-2 branch, gene discordance factor due to polyphyly (gDFP) = 65.37. NQS = normalized quartet score.
